# Oncogenic virus hijacks SOX18 pioneer function to enhance viral persistence

**DOI:** 10.1101/2025.06.28.662102

**Authors:** Krista Tuohinto, Matthew S. Graus, Peyton Staab, Ville Tiusanen, Fei Liangru, Susav Pradhan, Yew Yan Wong, Simon Weissmann, Jieqiong Lou, Elizabeth Hinde, Justin Wong, Quintin Lee, Alexey Terskikh, Martin Alvarez-Kuglen, Tara Karnezis, Thomas Günther, Adam Grundhoff, Biswajyoti Sahu, Mathias Francoís, Päivi M. Ojala

## Abstract

Kaposi’s sarcoma herpesvirus (KSHV) establishes lifelong oncogenic infection in lymphatic endothelial cells (LECs) by ensuring episomal maintenance of its genome via the viral protein LANA. Efficient viral genome maintenance typically involves host DNA replication and episome tethering, but the extent of cell-type-specific regulation remains unclear. Here, we identify that KSHV hijacks the pioneering function of the endothelial-specific transcription factor SOX18 to facilitate persistence of viral episomes. Upon infection, LANA co-opts SOX18 to recruit the SWI/SNF chromatin-remodeling complex via its ATPase subunit BRG1, enhancing chromatin accessibility and enabling efficient viral genome persistence. Disruption of SOX18 or BRG1—genetically or pharmacologically—leads to reduced episome load and attenuated hallmarks of virus infection. This work highlights how viruses can harness lineage-specific transcriptional regulators to establish persistent nuclear retention of their episome into the host genome.

## Introduction

Kaposi’s sarcoma (KS) is caused by an oncogenic human gamma herpesvirus (KSHV/HHV-8), during periods of immune deficiencies such as organ transplantations and among the 40 million people living with HIV worldwide. KS is considered to originate from endothelial cells and consists of excessive malformed vasculature with KSHV-infected spindle-shaped cells as a pathological hallmark (Gramolelli & Ojala, 2017). Due to the paucity of validated molecular targets, and a limited understanding of the molecular mechanisms that underpin long-term persistence, there is a clear limitation of current KS therapies, of which none are curative. Clinical outcomes are unfavorable especially in the lower income countries with the highest KS burden, such as sub-equatorial Africa, where it remains life-threatening for patients resistant to advanced anti-retroviral therapy (ART) (Cesarman et al., 2019).

We and others have shown that lymphatic endothelial cells (LEC) are exceptionally susceptible to KSHV infection in striking contrast to other cell types (Choi et al., 2020; DiMaio et al., 2020; Golas et al., 2019; Gramolelli et al., 2020). KSHV can reprogram blood vascular endothelial cells (BECs) toward a lymphatic endothelial-like state, driving a Kaposi’s sarcoma (KS) gene signature that more closely resembles LECs than BECs (Aguilar et al., 2012; Carroll et al., 2004; Hong et al., 2004; Wang et al., 2004). Remarkably, when KSHV infects LECs, the resulting KLECs undergo profound changes in morphology, proliferation, and identity—acquiring features characteristic of KS spindle cells (Cheng et al., 2011; Gasperini et al., 2012; Ojala & Schulz, 2014). However, the stepwise progression of the virus-driven molecular mechanisms that underpin this cell transformation are not well understood.

Latency is considered the default, quiescent mode of infection, where the circularized dsDNA KSHV genome persists as an extrachromosomal episome attached to host chromatin. A switch to lytic replication, triggered by the viral ORF50/RTA gene leads to the expression of all viral genes, resulting in the production and release of new virus particles (Han et al., 2024). Persistence of KSHV episomes is crucial for the establishment of a lifelong infection and for clinically evident KS to develop. KSHV Latency-Associated Nuclear Antigen (LANA) is required for tethering the viral episomes to host chromatin and for enabling KSHV to exploit the host replication machinery to replicate and maintain viral DNA in latently infected host cells (Ballestas & Kaye, 2001; Purushothaman et al., 2016; Uppal et al., 2014). LANA binds to the conserved terminal repeat (TR) sequences on the KSHV genome through its C-terminal domain and docks onto the host chromatin via histones H2A/B through its N-terminal domain (Barbera et al., 2006). Thus, LANA is required for viral episome tethering, latent replication and maintenance of the replicated genomes in daughter cells during mitosis. While LANA has been shown to generally bind to open or active chromatin (Hu et al., 2014; Kumar et al., 2022; Lotke et al., 2020; Lu et al., 2012; Mercier et al., 2014; Ye et al., 2024), how LANA may select for specific chromatin regions and which protein complexes are involved in stabilizing viral genomes remains a major question in the field.

KLECs display a unique infection program characterized by high numbers of intracellular viral episomes. In our previous work, we showed that SOX18 and PROX1, two key developmental transcription factors (TFs) for LEC fate, are widely expressed in KS tumors and critical to support this unique KSHV infection program in LECs by two distinct mechanisms (Gramolelli et al., 2020). SOX18 binds to the KSHV origins of replication, and its expression increases viral genome copies, while PROX1 plays a role in the reactivation of the lytic cycle.

The SOX18 TF is essential for embryonic development. It belongs to the SOXF group, which plays a role in vascular development angiogenesis, wound healing and cancer metastasis (Downes & Koopman, 2001; Schock & LaBonne, 2020). It is also involved in LEC differentiation by co-regulating PROX1 with NR2F2, a key regulator of lymphatic cell identity (Aranguren et al., 2013; Duong et al., 2012; Francois et al., 2008; Srinivasan et al., 2010). Because of its role in solid cancer development, SOX18 has long been a target for drug development. Despite long-standing challenges in targeting TFs with small molecules, the SOX18 inhibitor, Sm4, was identified (Fontaine et al., 2017; Overman et al., 2017). Mechanistically, Sm4 exerts its effects by selectively disrupting SOX18 dimerization, thereby suppressing its transcriptional activity. Importantly, SOX18 genetic depletion or its chemical inhibition by specific inhibitors, Sm4 or the R(+) enantiomer of propranolol (Holm et al., 2025; Overman et al., 2019; Seebauer et al., 2022), dramatically decreased the number of intracellular KSHV genome copies in KLEC indicating that SOX18 contributes to the maintenance of the high viral episome copies in KLECs (Gramolelli et al., 2020). Moreover, Sm4 also significantly decreased the infected spindle cell phenotype (hallmark of KSHV infection) and relative KSHV genome copies *in vivo* (Tuohinto et al., 2023), suggesting SOX18 as an attractive therapeutic target for KS.

To uncover how SOX18 regulates KSHV episome maintenance, we combined genomics, proteomics, and quantitative molecular imaging in infected lymphatic (LECs) and venous (HUVECs) endothelial cells. Using genome-wide chromatin profiling (ATAC-seq), proximity proteomics (BioID), and high-resolution imaging platforms—including MIEL for epigenetic landscape mapping (Farhy et al., 2019), Number and Brightness (N&B; (Hinde et al., 2016), and single-molecule tracking (Chen et al., 2014; McCann et al., 2021)—we dissected the step wise progression of the early molecular mechanisms that drive viral persistence. This integrative approach reveals a novel interplay between LANA, SOX18, and the chromatin remodeler BRG1 that orchestrates episome persistence and sustains KSHV latency.

## Results

### SOX18 recruits SWI/SNF chromatin remodeling complex upon KSHV infection

Our recent finding that SOX18 acts as a central hub for controlling viral genome copies during KSHV infection in LECs (Gramolelli et al., 2020) prompted us to investigate the molecular mechanism by which a host TF contributes to the maintenance of high viral genome levels and viral persistence. We first investigated whether SOX18 transactivation activity would be responsible for an increase in genome copies by activating expression of viral genes. To this end, we transduced lentiviruses expressing wild-type (wt) SOX18, two SOX18 mutants: a transactivation deficient, dominant negative mutant (C240X) and a DNA-binding HMG box deletion mutant (HMGdel; Fig S1A), or a Cherry expressing mock control, into HeLa cells inherently lacking SOX18 expression (Fig S1B). These cells were then subsequently infected with rKSHV.219 and analyzed for expression of selected latent (LANA, vCyclin, and vFLIP) and lytic (RTA, K-bZIP, and K8.1) viral genes at 72h.p.i. Intriguingly, no significant effects on transcription of the selected viral genes were seen in KSHV-HeLa cells expressing wt or either of the SOX18 mutants (Fig S1C). To further validate this observation, we measured mRNA levels of the same viral genes in KLECs treated with the SOX18 inhibitor (Sm4) and DMSO, as a control. As shown in (Fig S1D), the SOX18 inhibitor had no significant effects on transcription of the viral genes in KLECs within 24 hours, suggesting that SOX18 does not support high numbers of viral episomes via direct activation of viral gene expression. To further investigate another alternative mode of action, we next opted for an unbiased, proteomics-based approach to uncover SOX18 protein partners in KSHV-infected cells.

For this, a proximity-dependent biotinylation screen coupled with mass spectrometry (BioID) was carried out, allowing comprehensive and unbiased identification of proteins in proximity of SOX18. To this end, we transduced a lentivirus expressing BirA*-fusion of SOX18 into KSHV-infected cancer cells (iSLK.219; (Myoung & Ganem, 2011) and uninfected, parental (SLK) cells to differentiate interactions specific for KSHV-infection. The detection of SOX18 as one of the most highly enriched protein in both conditions served as an internal positive control for successful pull-down efficiency (Fig 1A-B). The BioID screen revealed cellular SWI/SNF chromatin remodeling complex (CRC) proteins as high confidence SOX18 interactors in the KSHV-infected cells (Fig 1A), but not in the parental uninfected cell line (Fig 1B-C). Most of the top SOX18 interactors in infected cells are components of the same CRC complex (Fig 1C).

**Figure 1.**
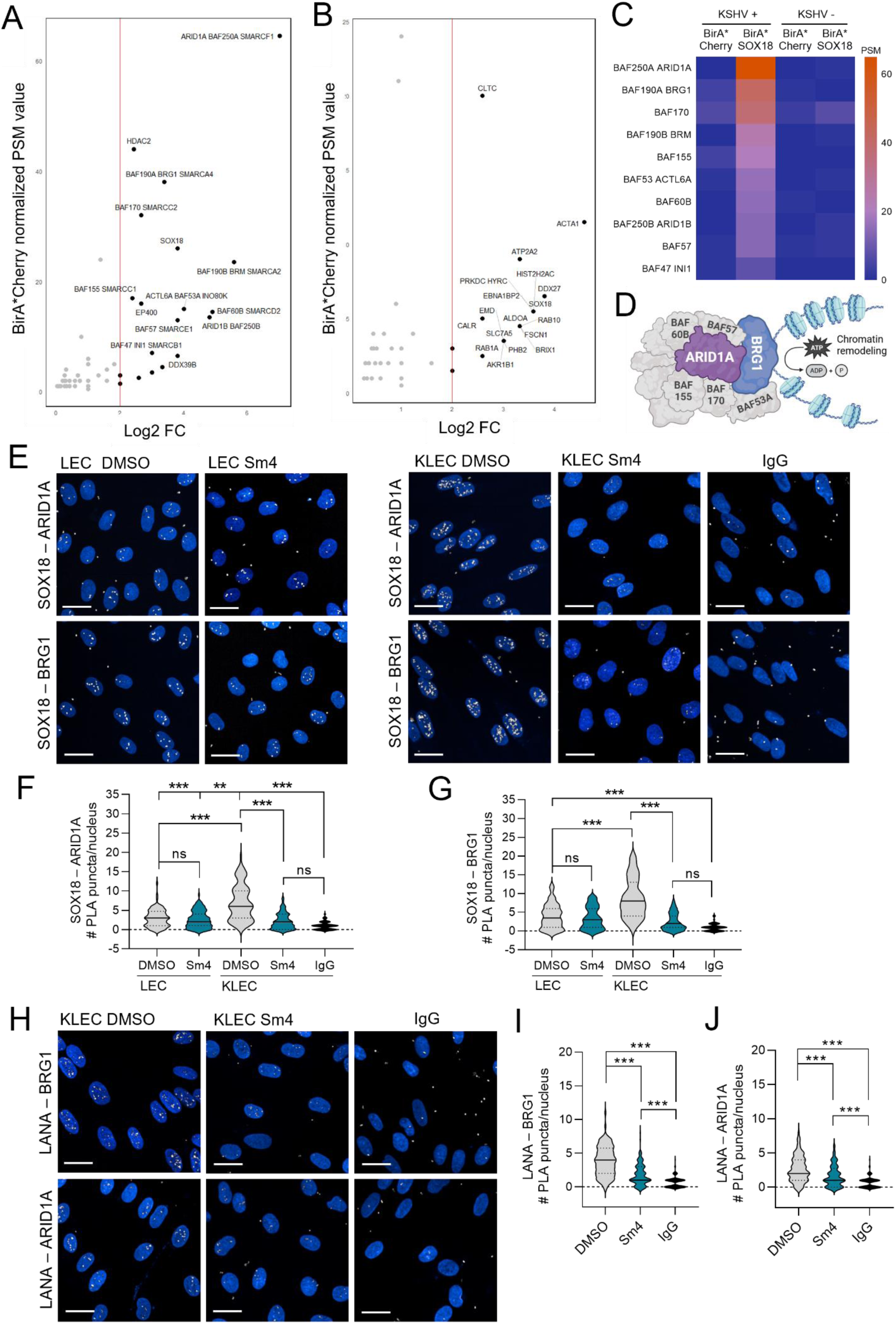
SOX18 recruits SWI/SNF chromatin remodeling complex upon KSHV infection. **A-B.** Bio-ID proximity-based protein-protein interaction screen using a BirA*-fusion of SOX18 (BirA*SOX18) or Cherry (BirA*Cherry) as a control in **A.** stably KSHV-infected iSLK.219 cells or **B.** parental, uninfected SLK cells. The strength of the interaction of SOX18 with the indicated proteins is shown as ≥ 2 log2 fold change FC (x-axis) and BirA*Cherry bait-normalized PSM = peptide spectral matches (y-axis). All shown proteins have ≥2 unique peptides. **C.** Heatmap of the canonical SWI/SNF (cBAF) complex subunits in all conditions. **D.** A schematic of the cBAF complex; ARID1A and BRG1, the top interactors of SOX18, are highlighted in purple and blue, respectively. **E-H.** Validation of the selected interactions by PLA with the indicated antibodies. **E.** PLA images of uninfected LECs (LEC) and rKSHV.219 -infected LECs (KLEC) at 72 h.p.i (hours post infection) treated with Sm4 or DMSO for 72h and imaged with Opera Phenix 40x, nuclei were counterstained with Hoechst (33342), scale bar is 20µm. **F-G.** Quantification of nuclear PLA puncta from 10 fields (n=100 nuclei) for **F.** SOX18-ARID1A and **G.** SOX18-BRG1 interactions in all conditions. **H.** LANA-BRG1 and LANA-ARID1A PLA images and **I-J.** quantification of nuclear PLA puncta from 10 fields (n=100 nuclei) in KLECs. Statistical significance was determined by ordinary one-way ANOVA with Tukey correction for multiple comparisons; *p < 0.05, **p < 0.01, ***p < 0.001, ns = non-significant.

SWI/SNF complex, also known as canonical BAF (cBAF), is a multi-subunit entity that confers ATPase activity to alter DNA-nucleosome contacts, thereby generating chromatin accessibility (Centore et al., 2020) (Fig 1D). SWI/SNF complex subunits BRG1 and ARID1A were among the top hits of SOX18 protein interaction partners in the infected cells. BRG1 is the catalytic ATPase remodelling the chromatin via nucleosome eviction and needed for efficient replication fork progression, whereas ARID1A serves as a stabilizing core of the complex, directing ATPase activity with high affinity to bind to chromatin and shown to interact with TFs (Cohen et al., 2010; Wanior et al., 2021).

To determine if ARID1A or BRG1 are modulated by KSHV infection or SOX18 inhibition in LECs, we confirmed the nuclear localization and protein expression levels in LECs and KLECs (Fig S1E-F). ARID1A levels were almost two-fold higher in KLECs over LECs and further increased with Sm4. Moreover, consistent with our previous findings, SOX18 levels were elevated following KSHV infection (Gramolelli et al., 2020). Next, we analyzed the interactions of SOX18 with BRG1 and ARID1A in LECs and KLECs treated with Sm4 using quantitative proximity ligation assay (PLA) image analysis (Fig 1E). In accordance with the BioID data, significantly higher number of PLA dots of SOX18-BRG1 and SOX18-ARID1A were seen in KLECs over uninfected LECs (Fig 1F-G). Further, uninfected cells treated with Sm4 showed negligible changes in the number of PLA puncta whereas in KLECs these interactions were significantly diminished by SOX18 pharmacological blockade without reducing their total protein levels (Fig 1F-G & Fig S1E-F).

As the interaction between SOX18 and SWI/SNF subunits was more pronounced in KLECs (Fig 1A, 1F-G), this led us to investigate whether a viral protein mediates these interactions. We recently demonstrated that SOX18 binds near the terminal repeat (TR) region of the KSHV genome (Gramolelli et al., 2020), which is also the binding site for LANA - a key initiator of latent viral DNA replication (Juillard et al., 2016; Schulz et al., 2023). Thus, we tested whether BRG1 or ARID1A also interact with LANA in KLECs. Interestingly, while LANA showed a clear interaction with BRG1 (Fig 1H, I), its interaction with ARID1A (Fig 1H, J) was only moderate in comparison to its interaction with SOX18 (Fig S1E, G). Notably, treatment with Sm4 significantly reduced the number of PLA dots, indicating that these interactions are SOX18-dependent. Collectively, these findings suggest that LANA facilitates SOX18 to recruit SWI/SNF primarily via BRG1 in KLECs. The discovery of a SOX18-SWI/SNF axis in KSHV-infected cells suggested a potential role for SOX18 as a virus-engaged pioneer factor. To test if KSHV infection enables SOX18 to function as a pioneer factor, we next set out to address whether SOX18 can influence chromatin organization independently of KSHV.

### Perturbations to SOX18 activity causes changes in chromatin accessibility

To address this, we used human umbilical vein endothelial cells (HUVECs) as a model system due to the endogenous expression of SOX18. We started by imaging HUVECs with two different techniques: 1) confocal microscopy and 2) stimulated emission depletion (STED). Imaging was performed on HUVECs treated with Sm4 and stained with SiR-DNA, a live-cell nuclear stain that preferentially intercalates into A-T rich regions, increasing fluorescent signal from heterochromatin due to its condensed form which can be seen as an increase in SiR-DNA signal (Fig 2A-B). STED imaging further shows at higher resolution the increased distribution and intensity of the heterochromatin (Fig 2A-C).

**Figure 2.**
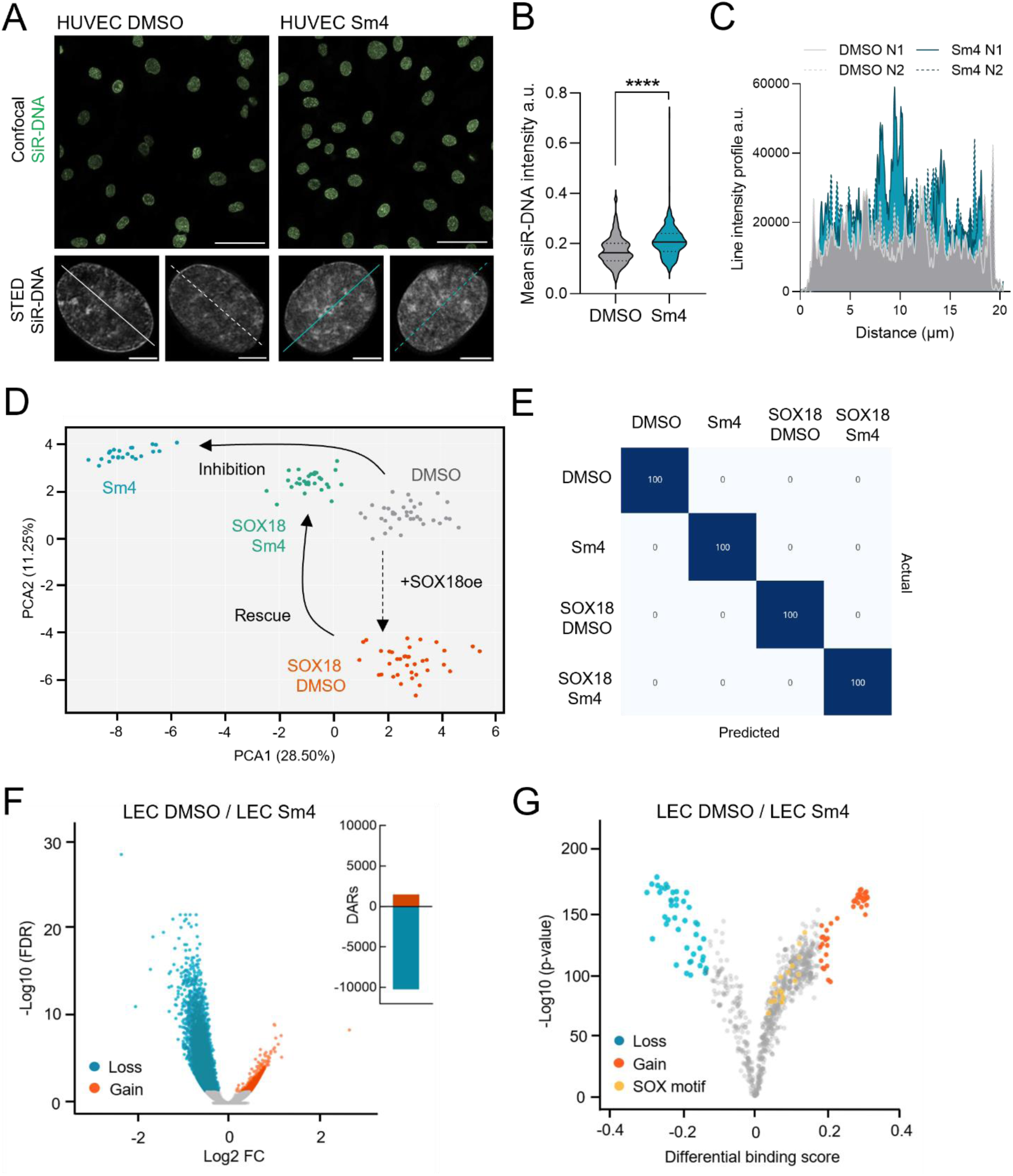
Perturbations to SOX18 causes changes in chromatin accessibility. **A.** Representative images of HUVECs treated with DMSO or Sm4 and stained with SiR-DNA, scale bar is 50μm for confocal images and 5μm for STED images. **B.** Mean intensity of SiR-DNA from cells in panel A, n ≥ 137 cells/condition. Statistical significance was determined by Mann-Whitney test ****p < 0.0001. **C.** Line intensity profile of STED images from panel A. Solid (N1) and dashed (N2) lines represent individual nuclei (N) line profiles. **D-E.** Quadratic discriminant MIEL analysis using texture features derived from images of HUVECs ± SOX18 over-expression and ± Sm4 treatment stained with DAPI. **D.** Scatter plot depict the first two discriminant factors for each cell population: each point is a pool of 60 cells. **E.** Matrix showing results for the discriminant analysis. Numbers represent the percent classified correctly (diagonal) and incorrectly (off the diagonal). **F-G.** LECs treated with DMSO or Sm4 for 24h and subjected for ATAC-seq. **F.** Data presented as volcano plot showing the human genomic regions with significant loss (turquoise) or gain (red) of accessibility and as number of differentially accessible regions (DAR) that have changed and **G.** as TF differential binding score volcano plot.

This observation is corroborated at the cell population level using a high-content, image-based, multi-parametric method known as microscopic imaging of epigenetic landscapes (MIEL; Farhy et al., 2019) (Fig S2A). Although we did not investigate epigenetic marks for these experiments, MIEL analysis is still able to measure differences in chromatin features on DAPI intensity across cell populations. To specifically evaluate if SOX18 dimerization influences chromatin organization, we examined HUVECs with ectopic over-expression of SOX18 to mimic the increase in SOX18 levels upon KSHV infection in LECs (Gramolelli et al., 2020). Similar to previous studies using MIEL analysis (Alvarez-Kuglen et al., 2024; Farhy et al., 2019), raw data was acquired from single cells (nuclei) of similar size and then pooled. The number of nuclei per pool was optimized to identify the minimum sample size yielding the maximal separation accuracy (n = 60 cells/point) (Fig S2B-D). As shown in Fig 2D MIEL analysis revealed significant changes in chromatin compaction upon SOX18 over-expression (oe; orange) whereas the opposite effect is observed after seven days of SOX18 inhibitor treatment (Sm4; blue), when compared to DMSO baseline control conditions (grey). In addition, gain-of-SOX18 function is rescued by its pharmacological inhibition, restoring a chromatin state close to the control conditions (green) (Fig 2D-E & Fig S2E). Importantly, TF-induced condensation is different from accessibility. Here, we suggest that SOX18, with its strong intrinsically disordered region (IDR; Fig S2F), may be driving chromatin into a more phase-separated, condensed state that still allows for transcriptional activity similarly to enhancer hubs or active transcriptional condensates (Boija et al., 2018; Erdos & Dosztanyi, 2024; Sabari et al., 2018).

To investigate whether SOX18 influences chromatin accessibility in LECs in the absence of KSHV infection, we performed Assay for Transposase-Accessible Chromatin-sequencing (ATAC-seq) on LECs treated for 24 hours with Sm4. This approach shows that pharmacological disruption of SOX18 activity significantly alters global LEC chromatin organization, when compared to DMSO treatment (Fig 2F), with over 10,000 differentially accessible regions (DAR) showing decreased accessibility (Fig 2F) and altered TF binding motifs in these DARs (Fig 2G). Global changes in chromatin accessibility can also be observed as a lower peak intensity in the heatmap showing the top 1000 sites with reduced accessibility (Fig S2G). Together, these observations demonstrate that SOX18 dimerization helps to maintain an open chromatin state, while its inhibition by Sm4 leads to a significant loss of chromatin accessibility.

### Chromatin compaction state feedback on SOX18 mobility and oligomeric states

To assess the interplay between different chromatin topological configurations and SOX18 biophysical behaviors, we combined two quantitative molecular imaging methods. Number and brightness (N&B; (Digman et al., 2008) analysis provides a map of SOX18 oligomeric states (Fig S3A), while single molecule tracking (SMT; (Chen et al., 2014) measures the mobility and chromatin binding dynamics of labeled molecules in real-time (Fig S3B). To perform these assays in live cells, we induced changes in the chromatin compaction state in HeLa cells using either Actinomycin D (ActD) to induce chromatin condensation, or Trichostatin A (TSA) to promote chromatin opening (Hinde et al., 2015; Hinde et al., 2016), while Halo-tagged SOX18 is ectopically expressed to enable live imaging at single molecule resolution.

Previous studies have shown that SOX18 dimer formation modulates an endothelial specific transcriptional signature (Moustaqil et al., 2018). The SOX18 protein dimerizes in a DNA-dependent manner via a cooperative binding mechanism on an inverted-repeat SOX motif spaced by five nucleotides (IR5), suggesting that molecular imaging should reveal at least a SOX18 population containing two states: monomers and dimers. Under baseline conditions (DMSO-treated, Fig 3A, top panels), N&B analysis revealed that SOX18 exists in a mixture of oligomeric states, with monomers being the most abundant, followed by dimers, which are relatively evenly distributed throughout the nucleus followed by the rare formation of higher-order oligomers, indicating that SOX18 can coexist in multiple multimeric states (Fig 3A-D). Chromatin opening with TSA treatment caused a significant increase in the formation of higher-order dimers and oligomers (Fig 3A, middle panels & Fig 3C-D), with monomers being found in between the higher-order collections. Conversely, induction of chromatin compaction with ActD led to an overall reduction of dimer formation and a higher-order oligomers formation with the whole population pre-dominantly found in a monomeric configuration (Fig 3A, bottom panels & Fig 3B-D). These observations are consistent with the fact that SOX18 dimer has low to no diffusion rates and further validate that its ability to self-assemble relies on template availability from the open chromatin (Moustaqil et al., 2018).

**Figure 3.**
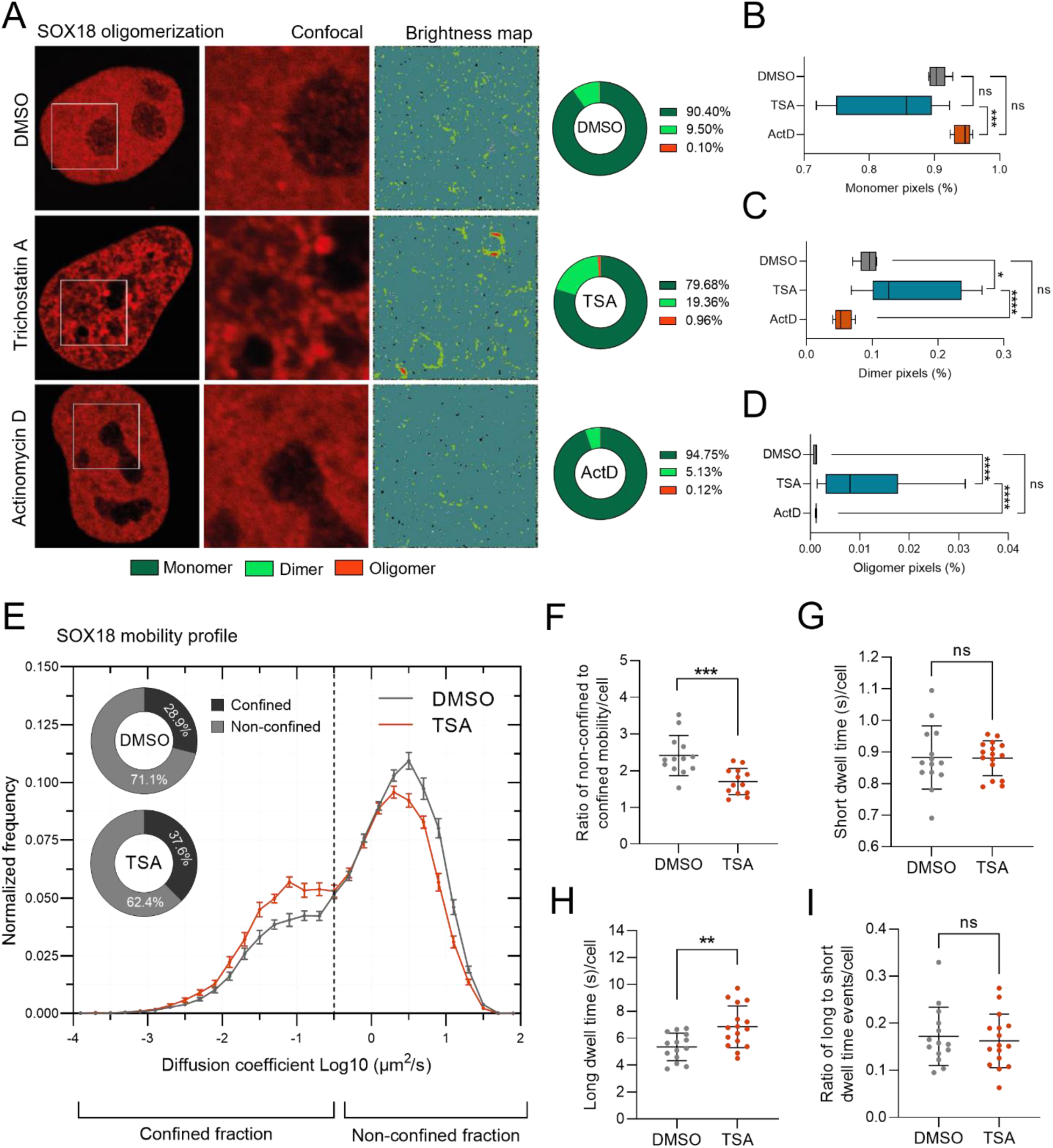
Chromatin compaction state feedback on SOX18 mobility and oligomeric states. **A.** Representative images and maps of SOX18 oligomeric states (monomer – dark green, dimer – light green, and oligomer – red) of HeLa cells transfected with SOX18 and treated with either DMSO, Trichostatin A (TSA), or Actinomycin D (ActD) and measured by N&B. **B-D.** Quantification of oligomeric states by N&B. For panels B-D, n > 5 cells, statistical significance was determined by by one-way ANOVA with a Dunnett correction for multiple comparisons, *p < 0.05, **p < 0.01, ****p < 0.0001. **E.** Diffusion mobility graph from single molecule tracking (SMT) acquisition comparing SOX18 mobility in HeLa cells that are transfected with Halo-SOX18 and treated either with DMSO or TSA. Pie charts represent the proportion of the trajectory population that is found in either the confined or non-confined states based on its diffusion coefficient. **F.** Ratio of non-confined to confined molecules per cell from F. **G-I.** Temporal occupancy characteristics for HeLa cells transfected with Halo-SOX18 and treated with TSA as G) short occupancy time, H) long occupancy time, I) ratio of long to short occupancy times. For panels G-I, n > 13 cells, statistical significance was determined by Welch’s t-test, **p < 0.01, ***p < 0.001.

To further assess the impact of chromatin accessibility on SOX18 behavior, we next measured its chromatin interaction dynamics under varying amounts of chromatin compaction using the SMT method. Here, we performed SMT on TSA treated cells and imaged SOX18 to assess its mobility profile and temporal occupancy (Chen et al., 2014; McCann et al., 2021). In all cells we can detect two populations of molecules based on their diffusion coefficient (Fig 3E). Cells treated with TSA showed a significant reduction in the diffusing population and an increase in the confined fraction compared to DMSO control (Fig 3E-F). These results parallel the N&B findings, which demonstrated an increase in SOX18 oligomerization in open chromatin conditions (TSA treatment; Fig 3A-D). We next questioned whether chromatin accessibility affects the SOX18-chromatin interaction lifetime. To test this, we measured the temporal occupancy of cells treated with TSA. The imaging determined that SOX18 short occupancy time, known as the target search mechanism for binding sites, remained unchanged compared to DMSO treatment (Fig 3G). By contrast, SOX18 long occupancy time, identified as a mechanism directly involved with transcriptional regulation per se, was significantly prolonged (Fig 3H). Additionally, the proportion of molecules that had prolonged interaction was not affected (Fig 3I). These results suggest that increased chromatin accessibility favours more stable SOX18-chromatin interactions involved with transcriptional modulation but not target genes search.

Taken together, the findings demonstrate that SOX18 forms higher-order oligomers (N&B) and that there is more SOX18 interacting with chromatin for a prolonged period of time (SMT) when more chromatin is accessible (TSA treated) (Table S1). These results then suggest that higher-order oligomers of SOX18 may have a more stable configuration, leading to increased duration of protein organizations. Studies in the p53 protein have also observed similar trends in high-order oligomers affecting its dissociation kinetics (Rajagopalan et al., 2011). Overall, the N&B and SMT analyzes reveal that SOX18 mobility, chromatin interaction dynamics and oligomeric states are directly dependent on open chromatin accessibility, supporting the notion that its pioneering function may play a role to promote its own activity as a transcriptional regulator.

Collectively, findings arising from a combination of genomics, super resolution and cell population-based quantitative imaging demonstrate that SOX18 activity significantly influences chromatin organization at multiple levels, via the modulation of mesoscale architecture and local regulatory regions accessibility.

### KSHV hijacks SOX18 pioneer activity to increase chromatin accessibility in LECs

The identification of a pioneer function for SOX18 in LECs prompted us to perform an ATAC-seq upon infection to investigate whether this role might be specifically hijacked by KSHV, as *de novo* infection increases SOX18 protein levels in LECs (Gramolelli et al., 2020). Since genomes of viral progeny resulting from spontaneous reactivation by rKSHV.219 in LECs would interfere with ATAC-seq analysis on viral genome, we opted to infect LECs with a strictly latent KSHV-BAC16 strain (ΔORF50; (Weissmann et al., 2025). Infection of LECs with ΔORF50 was first validated to induce the typical SOX18-dependent infection phenotypes as rKSHV.219 (see Materials and Methods & Fig S4) and confirming that the SOX18 upregulation and its critical functions occur during latency.

To investigate the link between SOX18-mediated chromatin changes and KSHV infection, we first sought out to delineate the effect KSHV has on chromatin organization by comparing ATAC-seq peaks of uninfected LECs to LECs infected with ΔORF50 (ΔORF50-KLEC). To ensure high quality data, we performed PCA and Pearson’s correlation analysis that indicated the robustness of the experimental reproducibility (Fig S5A-B). ATAC-seq analysis revealed that KSHV infection led to significant increases in chromatin accessibility on the host genome (Fig 4A, red), with over 22,000 regions becoming more accessible (red) and over 10,000 regions becoming less accessible (blue) when compared to uninfected LECs (Fig 4A, DARs: side panel). This indicates that KSHV infection alters host chromatin towards a more open state.

**Figure 4.**
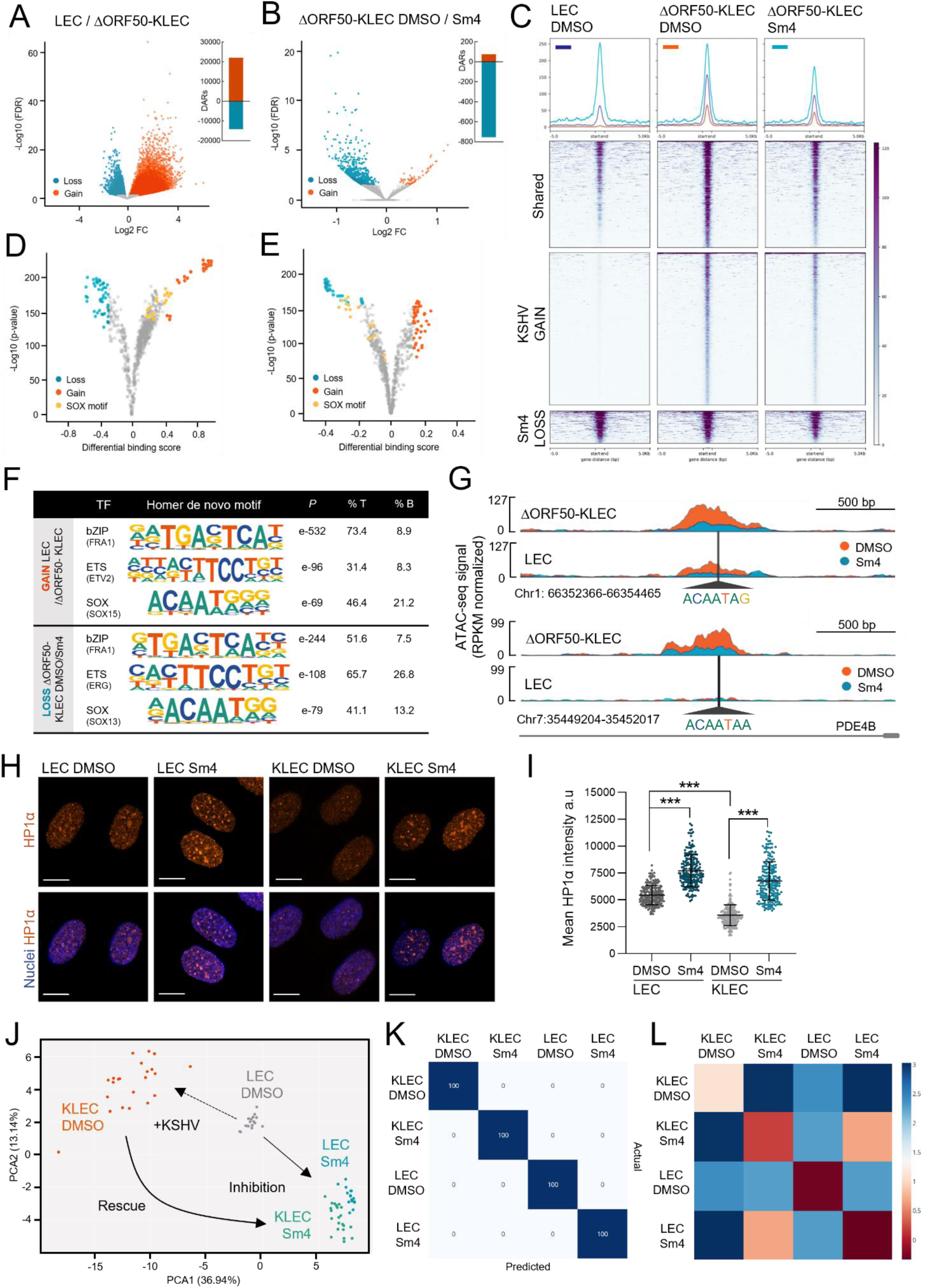
KSHV hijacks SOX18 pioneer activity to increase chromatin accessibility in LECs. **A-G.** Uninfected LECs (LEC) or LECs infected with KSHV-BAC16-ΔORF50 for 48h (ΔORF50-KLEC) were treated with DMSO or Sm4 for 24h and subjected for ATAC-seq. **A-B.** Volcano plots showing the human genomic regions with significant loss (turquoise) or gain (red) of accessibility upon A) KSHV infection and B) Sm4 treatment. **C.** Heatmap of the accessibility changes in the top 1000 sites upon KSHV infection (red line; KSHV gain), sites with accessibility loss after Sm4 treatment in ΔORF50-KLECs (light turquoise line; Sm4 loss) and sites with both accessibility gain upon infection and accessibility loss after Sm4 treatment (dark blue; shared). **D-E.** Volcano plots showing TF differential binding score prediction upon D) KSHV infection and E) Sm4 treatment in ΔORF50-KLECs. TFs with significant binding loss (turquoise), binding gain (red), and SOX family of TFs are marked (yellow). **F.** Top HOMER *de novo* transcription factor family motif enrichment gains upon infection and loss upon Sm4 treatment in ΔORF50-KLECs. **G.** Analysis of ATAC-seq peaks to show representative enhancer region with differential motifs and accessibility upon KSHV infection and SOX18 inhibition by Sm4 treatment in LECs and ΔORF50-KLECs. **H-I.** LECs infected with rKSHV.219 for 72h (KLEC) and treated with DMSO or Sm4 for 24h were H) labeled with anti-HP1α antibodies and imaged with Zeiss LSM880 confocal 63x for heterochromatin regions, nuclei were counterstained with DAPI, scale is bar 10µm. I) The mean nuclear intensity a.u. (arbitrary units) of HP1α signal quantified (n=200). Statistical significance was determined by one-way ANOVA with a Tukey correction for multiple comparisons, ***p < 0.001. **J-K.** Quadratic discriminant MIEL analysis using texture features derived from images of LECs and KLECs treated with DMSO or Sm4 24h and stained with DAPI and anti-HP1α antibodies. **J.** Scatter plot depict the first two discriminant factors for each cell population. Each point is a pool of 60 cells. **K.** Matrix showing results for the discriminant analysis. Numbers represent the percent classified correctly (diagonal) and incorrectly (off the diagonal). **L.** Average distance matrix calculated from the distance between each point per condition, with blue as farthest distances and red as closest distances.

With the understanding that KSHV infection increases chromatin accessibility and SOX18 inhibition reduces chromatin accessibility in uninfected LECs (Fig 2F-G & Fig S2G), we next assessed if the increased chromatin accessibility upon KSHV-infection is SOX18-dependent. To test this, we compared the ATAC-seq peaks upon DMSO and Sm4 treatment in ΔORF50-KLECs. ATAC-seq showed only minor changes on the viral chromatin itself (Fig S5C), further supporting that SOX18 does not contribute to high numbers of KSHV episomes in KLECs (Gramolelli et al., 2020) primarily by altering the transcription of viral genes (Fig S1C-D). However, chromatin accessibility of the host genome was significantly reduced when ΔORF50-KLECs were treated with Sm4 for 24 hours (Fig 4B), showing that inhibition of SOX18 is able to counter KSHV’s ability to induce increased chromatin accessibility. Importantly, the majority (76.7%) of chromatin regions that showed reduced accessibility with Sm4 were the same regions that were originally becoming accessible upon KSHV infection of LECs (Fig 4C, dark blue; shared).

We next examined the genomic regions that become more or less accessible due to infection or Sm4 treatment. Since we have determined that SOX18 dimerization contributes to chromatin organization, we asked whether SOX18 binding motifs become more accessible upon KSHV infection, which can lead to increased or sustained chromatin accessibility. The ATAC-seq data showed several SOX motifs becoming more accessible in infected LECs compared to the uninfected LECs (Fig 4D, yellow), which could be reversed upon Sm4 treatment (Fig 4E, yellow). We next determined through HOMER analysis that KSHV infection causes an enrichment for *de novo* motifs from bZIP (FRA1) and ETS (ETV2) TF families, in addition to SOX factors, while Sm4 treatment reciprocally showed reduced accessibility in the regions of corresponding TF family motifs (Fig 4F). This is further supported through KSHV-infection of LECs, which shows increased chromatin accessibility in enhancer regions (Fig 4G, red; DMSO), and SOX18 inhibition causing an opposite effect in the infected cells (Fig 4G, blue; Sm4). The enhancer regions also have an enrichment of the SOX18 motif. Here we use the variation in ATAC-seq peak height as a measure of chromatin accessibility to show that Sm4 treatment decreases accessibility (Fig 4G, red vs blue). These results further support the notion that SOX18 pioneer function plays a central role to enable KSHV-induced host chromatin accessibility in LECs.

We also assessed the chromatin compaction state by looking at HP1α, which is a known marker of heterochromatin formation and integrity (Schoelz & Riddle, 2022) as it can spontaneously phase-separate in solution, forming liquid-like droplets that preferentially sequester heterochromatin components such as nucleosomes and DNA, thereby promoting gene silencing (Bartkova et al., 2011; Larson et al., 2017; Strom et al., 2017). We did this by imaging LECs and KLECs treated with Sm4 and measuring HP1α intensity per cell and at the population level through MIEL analysis. Supporting the ATAC-seq findings, mean HP1α intensity was reduced upon KSHV-infection, whereas HP1α-associated heterochromatin foci were widespread and significantly more intense following Sm4 treatment both in LECs and KLECs (Fig 4H-I).

To assess whether SOX18 influences chromatin organization in LECs and KLECs, we performed MIEL as previously described in HUVECs (Fig 2D–E & S2A–E). In this experiment, we stained the cells with DAPI and an antibody for HP1α, comparing uninfected LECs to KLECs with and without Sm4 treatment (Fig S5D-H). As expected, KLECs (orange) have a distinctly different chromatin organization and HP1α distribution compared to uninfected LECs (grey). Additionally, we also find that the LEC and KLEC populations treated with Sm4 (respectively, blue and green) are more similar to each other than their respective DMSO controls (Fig 4J-L). An interesting observation from this experiment is the point distribution of LECs (grey) and KLECs (orange) that aligns with the Anna Karenina principle (Zaneveld et al., 2017), which posits healthy systems tending to be similar, while each dysfunctional system is abnormal in its own way. This is reflected in the broader distribution of chromatin features in DMSO treated KLECs where KSHV-infection is asynchronous compared to the more compact LEC populations (Fig 4L & Fig S5I). Interestingly, Sm4 treatment leads to a more compact chromatin feature distribution in KLECs, resembling the uninfected LEC state. This indicates that SOX18 inhibition can counteract the chromatin-altering effects of the asynchronous KSHV infection.

Taken together, the data demonstrates that KSHV hijacks SOX18 pioneer function and leads to a genome-wide change of host chromatin accessibility. To further investigate whether targeting the virus-induced host genome remodeling is a viable molecular strategy to reduce KSHV genome copies and hallmarks of infection, we next set out to target the SWI/SNF complex using either a pharmacological or a genetic interference approach.

### SWI/SNF ATPase activity is required for the hallmarks of KSHV infection in LECs

The observation of pharmacological blockade of SOX18 disrupting its interaction with ARID1A and BRG1 (Fig 1E-G) in KLECs prompted us to assess whether disruption of the SWI/SNF complex activity affects KSHV infection efficacy. To this end, we genetically depleted BRG1 and ARID1A by siRNA knockdown in KLECs (Fig 5A). Knockdown of either BRG1 or ARID1A resulted in a clear decrease in cell spindling (Fig 5B) and reduced the relative intracellular KSHV genome copies in KLECs (Fig 5C). Interestingly, depletion of BRG1 had a more pronounced impact on viral genome copy numbers compared to siARID1A, consistent with the stronger observed interaction between LANA and BRG1, than with ARID1A (Fig 1H-I & Fig S1G-H). Although significant, the reduction in KSHV genome copies was not as dramatic as seen with SOX18 inhibition by Sm4 (Fig S4D). This could be due to compensation by the ATPase BRM, also found among the SOX18 interactors by BioID (Fig 1A, C), and previously shown to be able to rescue BRG1 depletion (Hoffman et al., 2014).

**Figure 5.**
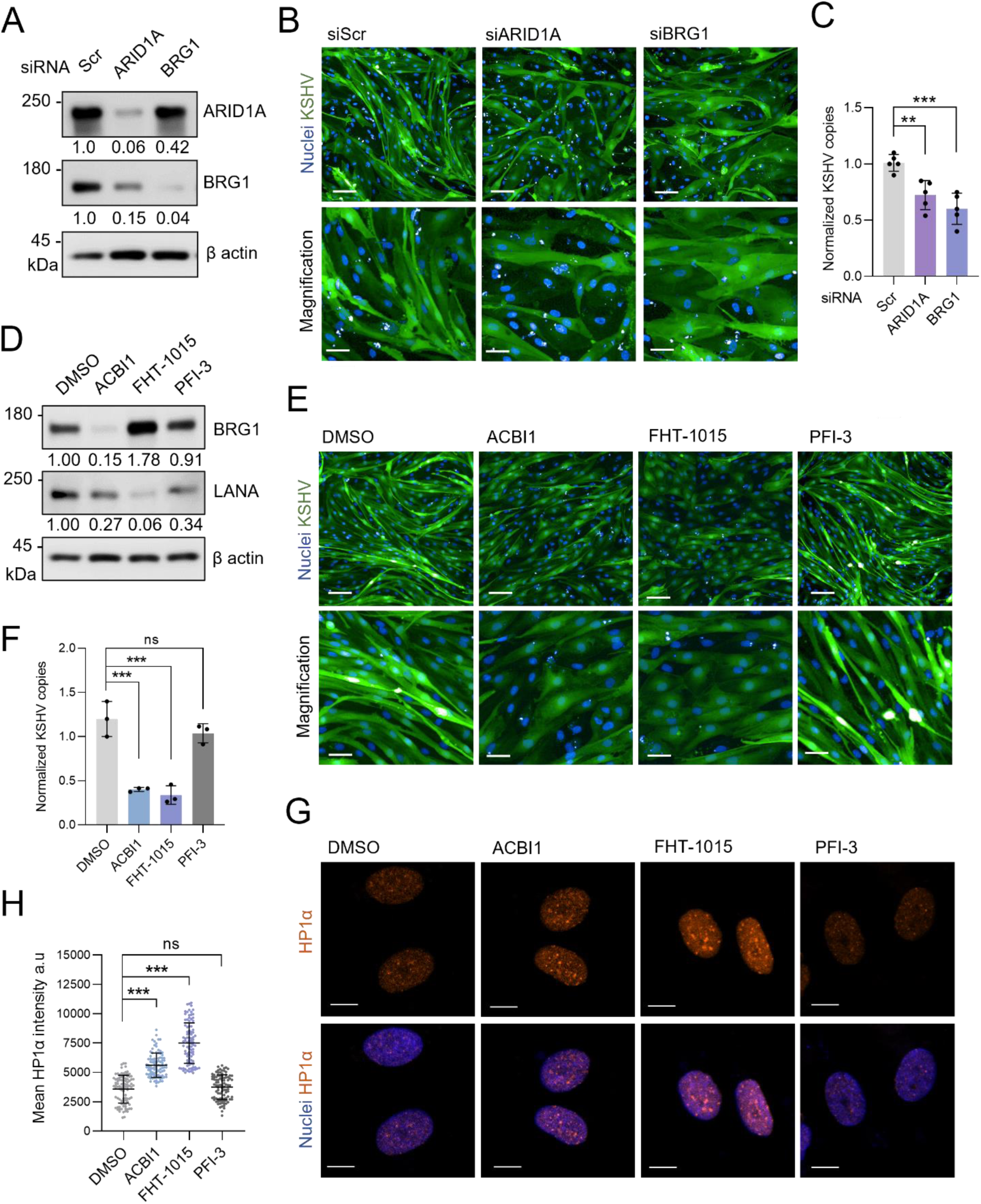
SWI/SNF ATPase activity is required for the hallmarks of KSHV infection in LECs. **A-C.** LECs were transfected with siRNAs targeting ARID1A, BRG1 or scramble (siScr) as a control for 24h, and thereafter infected with rKSHV.219 for 72h (KLEC). **A.** Immunoblotting for ARID1A and BRG1, and β-actin as a loading control. **B.** GFP-expressing KLECs imaged with Opera Phenix 20x for changes in the cell spindling phenotype. Nuclei were counterstained with Hoechst (33342), scale bar 100 µm, in magnification 30µm. **C.** KLECs quantified for normalized KSHV episome genome copies by qPCR (n=5). **D-H.** LECs infected with rKSHV.219 for 72h were treated with SWI/SNF inhibitors ACBI1, FHT-1015 and PFI-3 for 72h. **D.** Immunoblotting of KLECs for the indicated host proteins and LANA, and β-actin as a loading control. **E.** GFP-expressing KLECs imaged with Opera Phenix 20x for changes in spindling phenotype. Nuclei were counterstained with Hoechst (33342), scale bar is 100µm, in magnification 30µm. **F.** KLECs quantified for normalized KSHV genome copies by qPCR (n=3). **G.** Inhibitor-treated KLECs were labeled forHP1α and imaged with LSM 880 confocal 63x. Nuclei were counterstained with Hoechst (33342), scale bar is 10µm. **H.** The mean a.u. (arbitrary units) nuclear intensity of quantified HP1α signal (n=100 nuclei). Statistical significance was determined by one-way ANOVA with a Dunnett or Tukey correction for multiple comparisons, **p < 0.01, ***p < 0.001, ns = non-significant.

To avoid possible compensation by BRM, we obtained three specific inhibitors of the SWI/SNF complex. ACBI1 is a PROTAC (proteolysis-targeting chimera) inducing ubiquitylation and proteasomal degradation of the whole ATPase unit (BRG1/BRM), while FHT-1015 is an allosteric inhibitor of the BRG1/BRM ATPase (Battistello et al., 2023; Farnaby et al., 2019). PFI-3 is a selective bromodomain inhibitor for BRG1/BRM but does not detach the SWI/SNF from chromatin nor inhibit its ATPase activity (Singh et al., 2023; Wanior et al., 2021). A CTG cell viability assay using increasing concentrations of the inhibitors in LECs and KLECs showed a clear sensitization of KLECs to the BRG1/BRM ATPase inhibitors ACBI1 and FHT-1015, but not to PFI-3, when compared to uninfected LECs (Fig S6A-C). Degradation of BRG1 with the PROTAC ACBI1 was confirmed by Western blot, while the allosteric inhibitor FHT-1015 led to accumulation of BRG1 in the cells (Fig 5D). Moreover, relatively low concentrations of ACBI1 and FHT-1015 (30 nM and 10 nM, respectively) reduced the spindling phenotype of KLECs that showed reversal to the normal cobblestone EC morphology, while even high concentrations of PFI-3 did not (Fig 5E). Importantly, only inhibitors of the ATPase activity significantly decreased the KSHV genome copy numbers (Fig 5F), without affecting proliferation rates of treated cells (Fig S6D). In accordance with the significant reduction of relative KSHV episomes, LANA protein levels were decreased in cells treated with the ATPase inhibitors for 72 hours. Interestingly, PFI-3 moderately reduced LANA levels without significantly affecting the KSHV episome numbers. As siARID1A having a milder effect on infection than siBRG1 was already observed (Fig 5A-C), the inhibitor results further highlight the importance of the BRG1/BRM ATPase activity for KSHV infection over the core subunit ARID1A.

Immunofluorescence staining of HP1α in DMSO treated KLECs shows weaker signal when compared to either of the BRG1 ATPase inhibitor treated cells (Fig 5G). HP1α-associated heterochromatin foci were significantly more intense following ACBI1 and FHT-1015 but not PFI-3 treatment (Fig 5H), further supporting BRG1 ATPase activity involvement in SOX18 pioneer function in KLEC. The similar effects on heterochromatin upon Sm4 (Fig 4H-I) and FHT-1015 treatments prompted us to further compare the chromatin landscape following BRG1 or SOX18 blockade in KLECs. To this end, we performed ATAC-seq on KLECs following Sm4 or FHT-1015 inhibitor treatment (Fig S6E-F) and observed a striking overlap (61.5%) between the shared regions showing reduced accessibility upon SOX18 or BRG1 inhibition (Fig S6E, shared loss: dark blue peak).

These results demonstrate that KSHV instructs SOX18 to form a complex with the host SWI/SNF CRC and exploit its ATPase activity to alter host chromatin architecture and thereby maintain hallmarks of KSHV infection and high genome copy numbers in KLECs. To validate further the functional effects of SOX18 blockade on viral episomes, we next set out to disturb SOX18 activity.

### Effective KSHV episome maintenance relies on a functional SOX18

To uncover the molecular mode of action of SOX18 regulating the KSHV episome copies, we next assessed the effect of wtSOX18 and its C240X and HMGdel mutants on relative intracellular KSHV episome numbers. KSHV-HeLa cells transduced with mock (Cherry), wt and mutant SOX18-expressing lentiviruses prior to KSHV infection showed that only cells with wt, but not the mutants, contained a significantly higher relative number of intracellular episomes compared to Cherry control (Fig 6A). Moreover, confocal microscopy revealed that wtSOX18 expressing cells had the highest number of characteristic nuclear LANA speckles, which can be used as a surrogate marker for the viral episome numbers (Adang et al., 2006) (Fig S7A).

**Figure 6.**
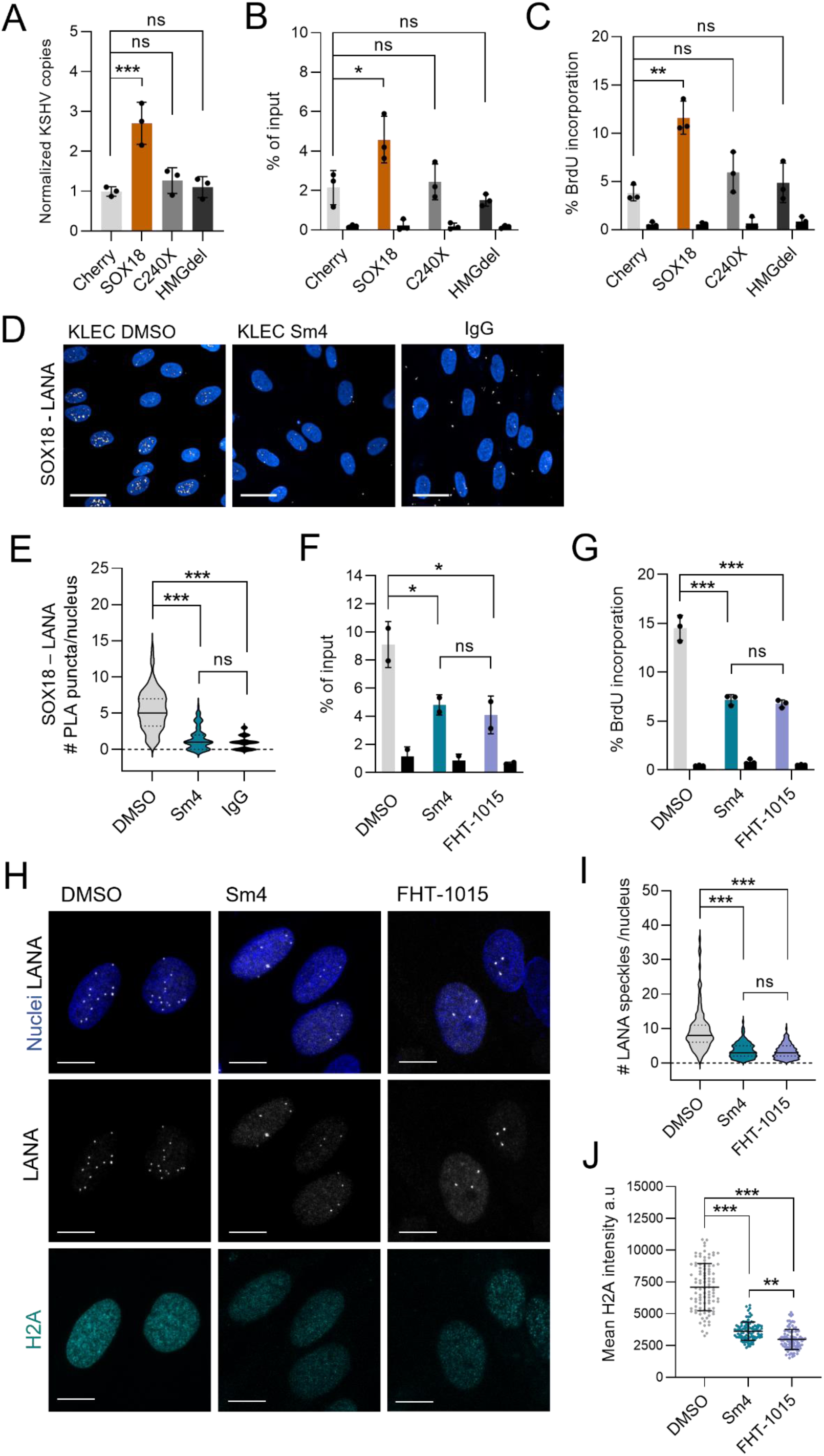
KSHV episome maintenance relies on a functional SOX18-BRG1 axis to increase LANA occupancy to KSHV TR. **A-C.** HeLa cells expressing SOX18wt, mutants C240X (dominant negative transactivation deficient) or HMGdel (DNA-binding deficient), or mCherry as a control, and thereafter infected with rKSHV.219 for 72h and A) measured for normalized KSHV genome copies by qPCR, B) subjected to ChIP-PCR with anti-LANA and IgG antibodies and analyzed for LANA binding at TR (n=3) or C) treated with BrdU for 4h and subjected to BrdU pulldown assay for nascent KSHV genome synthesis (n=3). **D-E.** KLECs treated with DMSO or Sm4 subjected to PLA assay using anti-SOX18 and anti-LANA antibodies, nuclei were counterstained with Hoechst (33342), D) imaged with Opera Phenix 40x, scale bar is 20µm and E) quantified as number of nuclear (n=100) PLA puncta (right panel). **F.** KLECs treated with DMSO, Sm4 or FHT-1015 for 24h were subjected to ChIP-PCR as described in B. (n=2). Two TR primers were used, and mean was taken for each replicate. **G.** KLECs treated with indicated inhibitors for 72h were subjected to BrdU pulldown assay as described in C (n=3). **H.** KLECs stained with anti-LANA and anti-H2A antibodies and imaged with LSM880 63x confocal, nuclei were counterstained with Hoechst (33342), scale bar is 10µm. **I-J.** Quantification of mean nuclear number of LANA speckles (n=100) and mean a.u. (arbitrary units) of H2A signal intensity (n=100). Statistical significance was determined by one-way ANOVA with Dunnett or Tukey correction for multiple comparisons, *p < 0.05, **p < 0.01, ***p < 0.001, ns = non-significant.

Our recent study demonstrated that SOX18 binds to DNA sequences within the terminal repeats (TR) of the viral episome (Gramolelli et al., 2020). Since LANA also binds to the TR, we next addressed if SOX18 contributes to LANA binding to the TR. Cherry, wtSOX18, or its mutant expressing KSHV-HeLa cells were subjected to LANA ChIP-PCR using primers for known LANA binding sites (LBS) on the TR. LANA ChIP-qPCR showed that LANA occupancy to TR was significantly increased in cells expressing wtSOX18 when compared to the mutants or the Cherry control (Fig 6B). We then sought to investigate if SOX18, as a secondary effect, would promote latent viral DNA replication directed by LANA from the TR origin of replication. To this end, we subjected the transduced KSHV-HeLa cells to a BrdU-pulldown assay. Only wtSOX18, but neither of the mutants, could significantly increase the newly synthesized, nascent viral DNA when compared to Cherry control (Fig 6C). KSHV, as other DNA viruses, utilize the host cell replication machinery to replicate their latent DNA genome once during the cell cycle. We therefore performed EdU cell proliferation assay and confirmed that the increases in KSHV episomes and DNA synthesis were not simply due to increased proliferation of wtSOX18-expressing cells (Fig S7B). These findings suggest that SOX18 needs both its intact transactivation and DNA binding domains to increase the occupancy of LANA to TR, thereby leading to more efficient latent viral DNA synthesis and increase in viral episome copies.

The correlation between SOX18 with increased LANA occupancy to TR prompted us to address if SOX18 itself interacts with LANA in KLECs. Interaction of SOX18 with LANA was confirmed by PLA and abolished by Sm4 treatment (Fig 6D-E). These findings indicate that SOX18 binds not only to the TR (Gramolelli et al., 2020) but also to LANA leading to higher occupancy of LANA on the KSHV origin of latent replication. Since we identified a critical SOX18-SWI/SNF axis on maintaining the high episome numbers, we next assessed the functional role of BRG1 in the LANA-SOX18 complex.

### BRG1 activity increases LANA occupancy at TRs in a SOX18-dependent manner

Since LANA interacts with both SOX18 (Fig 6D–E) and BRG1 (Fig 1H–I) in KLECs, we first investigated the contribution of BRG1 to the SOX18-dependent increase in the binding of LANA to TR in KLECs as described in (Fig 6B). To confirm specificity, we chose to add two primers for non-LANA binding sites on KSHV genome as well as two additional primers for human genome as negative controls. As shown by ChIP-QPCR in (Fig 6F), both inhibitors reduced the occupancy of LANA to TR within 24 hours but had no significant effects on the negative control sites on viral or host genome (Fig S7C). Immunoblotting confirmed that the reduced LANA binding was not due to lower levels of LANA protein in the inhibitor treated cells (Fig S7D-E), indicating that both inhibitors can specifically reduce LANA binding at TR. We then again performed the BrdU-pulldown assay to investigate the effects of Sm4 and FHT-1015 on nascent viral DNA synthesis, which showed a significantly reduced viral DNA synthesis rate following inhibitor treatments when compared to the DMSO control (Fig 6G). In conclusion, gain of SOX18 in KSHV-HeLa and in KLECs increases LANA occupancy at TR as a primary effect, and consequently leads to higher viral DNA synthesis rate and increased number of intracellular episomes. Both LANA binding to TR and latent viral DNA synthesis can be significantly diminished by chemical blockade of the SOX18-BRG1 axis.

Importantly, upon treatment with both inhibitors the number of LANA speckles originating from LANA clustering and formation of higher-order oligomers decreased at TRs, (Hellert et al., 2015) with the signal of LANA pattern appearing less dense (Fig 6H, top and middle panels, Fig 6I). This indicates a more diffusive behaviour of LANA with reduced binding to KSHV episomes. To address whether the SOX18-BRG1 axis would contribute to LANA-mediated KSHV episome tethering onto host chromatin via histone H2A and H2B (Ballestas & Kaye, 2001; Barbera et al., 2006; Verma et al., 2013), we analyzed histone expression upon Sm4 or FHT-1015 treatment of KLEC. IF analysis revealed a significant concurrent reduction in LANA speckles and H2A intensity in inhibitor-treated KLECs (Fig 6H-J). Reduction in both H2A and H2B was further confirmed by immunoblotting in Sm4 and FHT-1015 treated KLECs (Fig S7F).

These findings reveal that the SOX18–BRG1 axis contributes to the high KSHV episome numbers in KLECs by initially promoting host chromatin reorganization and LANA occupancy to TR, thereby potentially facilitating more efficient LANA-mediated replication and tethering of viral episomes to host chromatin, which ensures persistent and unique infection phenotype in KLECs.

## Discussion

Beyond transcriptional regulation, TFs are increasingly recognized as key regulators of chromatin organization and condensation state, processes that are essential for homeostasis, differentiation, and cell fate (Shaban et al., 2024). TFs have been classified in three classes: pioneer (control chromatin accessibility), settler (maintain chromatin conformation), and migrant (modulate transcription rates) (Sherwood et al., 2014). Pioneering activity is a unique ability of some TFs to bind directly to condensed chromatin to initiate chromatin remodeling events and increase chromatin accessibility (Bulyk et al., 2023). While direct evidence of viruses specifically hijacking pioneer transcription factors is limited, there are notable examples where viruses interact with host transcription machinery in a manner reminiscent of pioneer factor activity (HBV/HNF4a and HPV/E2) (Neugebauer et al., 2023).

Our study reveals how the endotheliotropic, oncogenic KSHV exploits two previously unrecognized aspects of SOX18 biology to promote viral persistence in LECs (Fig 7). First, SOX18 exhibits pioneer transcription factor activity by recruiting the SWI/SNF complex, thereby altering the host chromatin architecture by modulating other TFs’ accessibility. Second, through its interaction with the multifunctional, viral protein LANA, SOX18 facilitates the docking and stabilization of viral episomes onto the remodeled host chromatin, acting as a settler-like TF. These newly identified functions of SOX18 underscore its pivotal role in viral infection and provide a mechanistic basis for the virus-induced upregulation of SOX18 at both mRNA and protein levels in infected endothelial cells.

**Figure 7.**
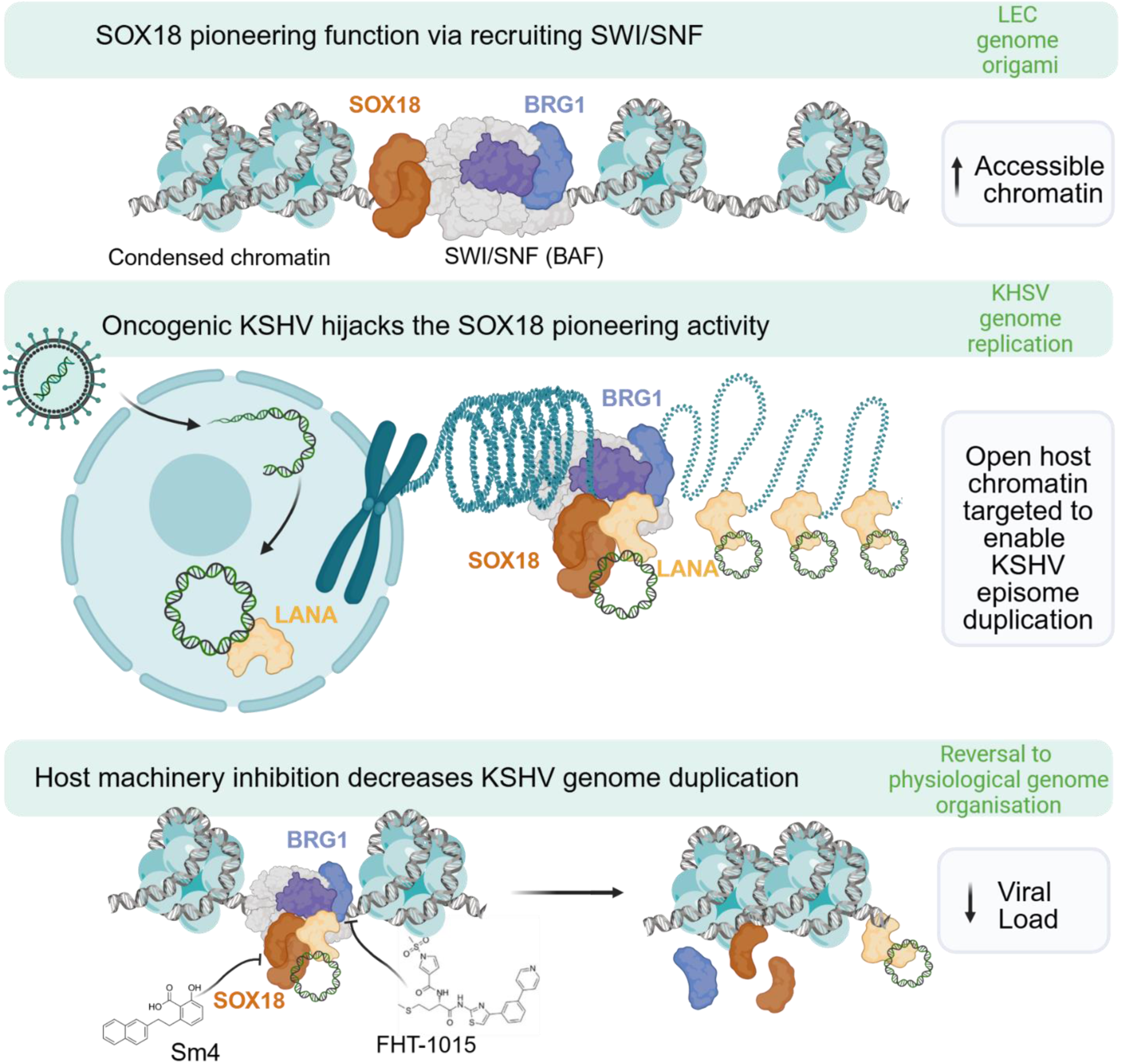
Graphical abstract. *Top panel:* The SOX18 transcription factor exhibits a pioneering role through its interaction with the SWI/SNF complex, shaping chromatin accessibility and genome organization in LECs. *Middle panel:* Upon KSHV infection, SOX18 is upregulated, and the viral LANA protein hijacks the SOX18/BRG1 pioneer complex to anchor viral episomes onto the host genome. This LANA–SOX18– BRG1 axis establishes a chromatin environment conducive to latent viral genome replication. *Bottom panel:* Pharmacological disruption of the host chromatin machinery impairs SOX18 or BRG1 function and consequently inhibits viral genome duplication. This highlights previously unrecognized host-derived therapeutic targets for the treatment of KSHV infection and the associated diseases.

A classic example of a TF with pioneer function is SOX2, which maintains the pluripotency of embryonic stem cells through its pioneer activity (Hagey et al., 2022). Despite their disparate roles, previous works have indicated that certain SOX factors, including SOX18, can replace SOX2 during stem cell reprogramming, albeit with a significantly lower efficiency (Nakagawa et al., 2008). The structure of SOX2 HMG domain, needed for pioneer DNA binding, was recently shown to be conserved among several SOX factors, including SOX18 (Dodonova et al., 2020). Previous studies have found SOX18 expression in stromal-derived adipose cells can induce endothelial cell markers, such as PECAM1, VE-Cadherin, KDR, and CD34, or induce hemogenic endothelium derived progenitors towards NK lymphoid pathways, demonstrating SOX18’s ability to reprogram cell identity (Fontijn et al., 2014; Jung et al., 2023). Our discovery of SOX18 pioneer function now enforces the capacity of SOX18 to exert pioneering activity and reprogramming, in a context-dependent manner.

In pathological conditions, such as cancer or infection, otherwise non-pioneering TFs at physiological levels can gain pioneer activity due to mutation or overexpression (Bulyk et al., 2023). This virus-instructed SOX18 pioneering activity also likely contributes to the KSHV-driven reprogramming of ECs, as reported in our prior studies and those of others (Aguilar et al., 2012; Carroll et al., 2004; Cheng et al., 2011; Gasperini et al., 2012; Hong et al., 2004; Wang et al., 2004). Here we demonstrate that SOX18 does not directly regulate viral gene expression but is redirected by KSHV to remodel the host chromatin architecture. This provides a novel perspective on viral infection, showing that a host transcriptional regulator can be hijacked to perform alternative molecular roles that favor viral DNA replication and genome maintenance. Notably, Epstein Barr virus (EBV), a close gamma herpesvirus family member of KSHV, has also been found to induce reprogramming through chromatin accessibility (Ka-Yue Chow et al., 2022), albeit the molecular mechanisms that drive these changes of the epigenome landscape remain unknown.

The identified LANA–SOX18–BRG1 axis promotes a permissive chromatin environment, essential for LANA binding to the TRs on viral episome, efficient latent viral DNA replication and possibly also episome docking to the host genome. Our findings are nicely corroborated by a previous report demonstrating an interaction between SWI/SNF complex subunits and LANA (Zhang et al., 2016), as well as BRG1 and TR (Si et al., 2006). The importance of the SWI/SNF complex in KSHV pathobiology is further supported by its previously described role in RTA/ORF50-mediated viral lytic replication (Gwack et al., 2003; Lu et al., 2003). However, this study demonstrates that the interaction of SOX18 with the SWI/SNF complex and the phenotypic changes upon SOX18 or BRG1 inhibition in KLECs occur during latency.

Our work suggests a new approach to target oncogenic viral infection by demonstrating that targeting host transcriptional modulators that directly engage with KSHV are viable molecular targets. This redefines the current dogma for anti-viral therapies that mostly relies on targeting viral effectors rather than host targets. As critical regulators of gene expression and key drivers of cancer and other diseases, components of the SWI/SNF chromatin remodeling complex have emerged as promising therapeutic targets. Pharmacological inhibition of SWI/SNF, particularly through agents targeting its catalytic subunits or via proteolysis-targeting chimeras (PROTACs), has demonstrated substantial therapeutic potential in extensive preclinical studies across a wide range of cancer types (Battistello et al., 2023; Centore et al., 2020; Farnaby et al., 2019). Importantly, some of these agents have progressed from preclinical research to clinical trials, showing promise of developing effective cancer therapeutics targeting the SWI/SNF complex functions (Dreier et al., 2024; Malone & Roberts, 2024). In our KLEC model, only ATPase inhibitors targeting BRG1 were effective in reducing key hallmarks of infection, whereas inhibition of the BRG1 bromodomain by PFI-3 had no observable effect. This suggests that while PFI-3 may disrupt the recruitment or stabilization of SWI/SNF complex at chromatin, thereby impairing remodeling activity (Lee et al., 2021; Wanior et al., 2021), it may not inhibit the ATPase function of BRG1 once it is already recruited to chromatin by LANA and SOX18.

Our previous findings suggested SOX18 as an attractive therapeutic target for KS (Gramolelli et al., 2020; Tuohinto et al., 2023) and that this TF activity is directly modulated as part of R(+) -Propranolol off-target effects (Holm et al., 2025; Overman et al., 2019; Seebauer et al., 2022) in the context of infantile hemangioma. This is further endorsed by a recent study where a 6-month oral propranolol treatment of a patient with classic KS resulted in a substantial decrease in the size of skin KS lesions associated with a reduction in KSHV infection (Salido-Vallejo et al., 2022). We further identify the SOX18–BRG1 axis as a key regulator of viral latency maintenance in LEC with important implications for the development of therapeutic strategies targeting chromatin regulators as a potential molecular approach for managing both KSHV infection and Kaposi Sarcoma.

### Limitations of the study

One of the limitations of our study is that we do not confirm all the findings in LECs. We used HUVECs and HeLa cells as model systems to investigate how SOX18 navigates the nuclear environment. Specifically, how pharmacological inhibition of SOX18 changes the chromatin organization (HUVECs) and, in turn, how chromatin organization alters SOX18 genome navigation in real-time (HeLa). In venous HUVECs, SOX18 is expressed to differentiate towards lymphatic endothelial lineage, whereas LECs are considered the KS cell of origin and display a unique KSHV infection program with SOX18 upregulation. Nevertheless, combining unbiased proteomics screen (BioID), genomics-approach (ATAC-seq), and large-scale chromatin imaging (MIEL), we have discovered SOX18 pioneer function in both ECs independently, in physiological and pathological conditions, and via various approaches that all support SOX18 pioneer function.

Another limitation of our current study is that our results only provide circumstantial evidence that SOX18 increases KSHV latent DNA replication and indicate that SOX18 is required to alter the host chromatin for efficient episome stabilization and thereby support higher viral DNA synthesis rates by ensuring higher numbers of template episome genomes. Our aim here was to define the mechanism of how SOX18 promotes high numbers of viral episomes, to pave way for future translational studies where the efficacy of inhibiting chromatin remodelers can be assessed. Therefore, we did not test the efficacy of BRG1 ATPase inhibitors or PROTACs in a preclinical KS model as this was not in the scope of this study.

## Acknowledgements

We acknowledge and are extremely grateful to Endrit Elbasani, Silvia Gramolelli, Veijo Nurminen, Lorenza Cutrone, and Riikka Kallinen for their valuable contributions to this study, and Nadezhda Zinovkina for technical assistance. We also acknowledge the technical staff of Biomedicum Imaging Unit (BIU) and Finnish Institute of Molecular Medicine (FIMM) for their help in IFA imaging, and Proteomics Unit (Institute of Biotechnology, University of Helsinki) in BioID processing and analysis. We thank Norwegian Sequencing Centre, Oslo, Norway, for sequencing services. We kindly thank Riikka Kivelä (University of Jyväskylä and RPU, University of Helsinki, Finland), Bala Chandran (USF, FL, USA), Carolina Arias (Biohub SF, CA, USA) for reagents.

We also acknowledge the technical and scientific staff of Sydney Microscopy & Microanalysis, the University of Sydney node of Microscopy Australia for their consultation and assistance with the STED, MIEL, and SMT experiments. The authors thank Dr. Rosemary Steinberg for her consultation and advice on statistical analysis and Dr. Kenta Ninomiya for his consultation and advice on MIEL analysis.

## Funding

PMO: Sigrid Juselius Foundation, Finnish Cancer Foundation, The Gertrude Biomedical Pty Ltd.

KT: Doctoral program in Biomedicine, University of Helsinki, Finnish Cultural Foundation, Biomedicum Foundation, Magnus Ehrnrooth foundation.

PS: ICANDOC doctoral pilot program.

MF: NHMRC Idea Grants (APP2019904 and APP2029719) and NIH grant 5R01HL096384.

MG: NIH grant 5R01HL096384.

BS: Finnish Cancer Foundation, Sigrid Jusélius Foundation, Jane and Aatos Erkko Foundation, iCAN Digital Precision Cancer Medicine Flagship (320185), Norwegian Cancer Society (274630), South-Eastern Norway Health Authority (2025083) and Norwegian Centre for Molecular Biosciences and Medicine, University of Oslo.

VT: Doctoral program in Integrative Life Sciences, University of Helsinki

SW, TG, AG: Deutsche Forschungsgemeinschaft (DFG) grant GR 3318/5-1 (Research Unit FOR5200 DEEP-DV, project number 443644894).

## Author contributions

KT, MG, MF and PMO conceived and designed the experiments with the help of SW, TG, AG, TK and BS. KT, MG, PS, LF, SW, SP and JQ performed the experiments. KT, MG, PS, VT, YW, EH, AT, and MK analyzed the data. QL and JW produced reagents for experiments. KT, MG, MF and PMO wrote the original draft of the manuscript. Manuscript editing and reviewing – KT, MG, PS, VT, SW, AG, BS, MF and PMO.

## Materials and methods

### Cell culture

Primary human dermal lymphatic endothelial cells LEC (Promocell; C-12216) were maintained in Microvascular MV-2 (Promocell; C-22121) medium supplemented with 5% fetal bovine serum, basic fibroblast growth factor, insulin-like growth factor 3, epidermal growth factor, gentamicin sulfate/amphotericin, ascorbic acid, and hydrocortisone; VEGF was not added. LECs were used until passage five. HUVEC cells (ATCC) were grown in EGM-2 media supplemented with EGM-2 bullet kit (Lonza; CC-3202).

iSLK.219 (Myoung & Ganem, 2011) is an RTA -inducible renal-cell carcinoma SLK cell line, stably infected with a recombinant KSHV.219. HeLa, SLK and iSLK.219 were grown in DMEM (BioNordika; ECB7501L), supplemented with 10% FBS (Gibco; 10270-106), 1% L-glutamate (BioNordika; ECB3000D), and 1% penicillin/ streptomycin (BioNordika; ECB3001D). iSLK.219 cells were also supplied with 10μg/mL puromycin (Sigma; P8833), 600μg/mL hygromycin B (Invitrogen; 687010), and 400μg/mL Geneticin G418 (Roche; 04727878001). iSLK.BAC16-ΔORF50 cells were supplemented with 0.5µg/ml puromycin, 200µg/ml hygromycin, and 1000µg/ml G418. Cell lines were used for approximately 15-20 passages.

All cells were propagated in a humified incubator at standard conditions. Cells were regularly tested negative for *Mycoplasma* (MycoAlert Mycoplasma Detection Kit, Lonza; LT07-705).

### Virus production and infections

Lentivirus production was performed as described in (Gramolelli et al., 2018). The concentrated virus preparation of recombinant KSHV.219 virus was produced from iSLK.219 (Myoung & Ganem, 2011) as described in (Tuohinto et al., 2023) and the virus was precipitated with PEG-it (Systems Biosciences; LV825A-1). Cells infected with rKSHV.219 express green fluorescent protein (GFP) from the constitutively active human elongation factor 1-a (EF-1a) promoter and red fluorescent protein (RFP) under the control of RTA-responsive polyadenylated nuclear (PAN) promoter, expressed only during lytic replication. An ORF50 deletion mutant KSHV-BAC16-ΔORF50 (KSHV-ΔORF50) virus was generated as described in (Weissmann et al., 2025) and produced and concentrated from iSLK.BAC16-ΔORF50 cells similarly as rKSHV.219. Cells infected with KSHV-ΔORF50 express green fluorescent protein (GFP) from the constitutively active human elongation factor 1-a (EF-1a) promoter. The concentrated virus was resuspended in ice-cold PBS, snap-frozen and stored at −80°C.

For experimental assays, cells were infected with low titers (MOI 1-2) of rKSHV.219, KSHV-ΔORF50, or transduced with lentiviruses in media with supplements in the presence of 8 μg/mL polybrene (Sigma; H9268) and spinoculation at 450 x g for 30 min, RT with the 5804R centrifuge (Eppendorf). Around 90-100% KSHV infection efficiency was achieved without selection. Virus titers were determined by infecting naïve LECs using serial dilutions of the concentrated virus and assessing the amount of GFP+ and LANA+ cells 72h post-infection with Phenix Opera 20x.

### Plasmid constructs

The BirA*SOX18 plasmid construct for BioID was generated from pFuW-myc-BirA-NLS-mCherry (a kind gift by R. Kivelä, University of Helsinki), used as a BirA*Cherry control. The wild-type human SOX18 insert sequence was codon optimized and synthesized by GeneArt (Thermo Fisher) to reduce G-C content, and pFuW-myc-BirA-NLS was inserted to N-terminus of SOX18. The resulting BirA*SOX18 consists of biotin-binding BirA, a nuclear localization signal (NLS), an SOX18 ORF (1152-5bp), including DNA-binding HMG domain (247-462bp), homodimerization domain (463-597bp), and transactivation domain (502-780bp).

A pFuW-myc backbone was also used to produce the Cherry, SOX18wt, and mutant plasmids C240X and HMGdel. The C240X and HMGdel mutants are described in (McCann et al., 2021). The codon optimized SOX18wt, C240X, and HMGdel sequences were then cloned in a pFuW-myc plasmid by Gibson Assembly. Both the backbone and gene inserts for SOX18wt, C240X, and HMGdel were assembled using NEB HiFi DNA assembly (NEB; E2611). Sanger Sequencing performed verification of the inserts while restriction analysis was performed to verify the integrity of the backbone.

### Inhibitor treatments

Small molecule SOX18 inhibitor Sm4 (Sigma; SML1999 / or a kind gift from Gertrude Biomedical Pty Ltd., Australia) was solubilized in DMSO (Sigma; D8418), stored in −80°C, and mixed with cell media at 20µM for LECs and KLECs and 30µM for HUVECs. ACBI1 (MedChemExpress; 128359) was solubilized in DMSO, stored in −80°C and mixed with cell media at 30nM. FHT-1015 (MedChemExpress; 144896) was solubilized in DMSO, stored in −80°C and mixed with cell media at 10nM. PFI-3 (Sigma; SML0939) was solubilized in DMSO, stored in −80°C and mixed with cell media at 50µM.

### Transfections

Transient transfection of siRNA of a semi-confluent culture of KSHV-infected LEC was done using OptiMEM (Gibco; 31985047), 1.5µl of Lipofectamine RNAiMAX (Invitrogen; 13778075) and 25nM siRNA per well in a 12-well plate according to manufacturer’s instructions with MV-2 media. Next day cells were supplied with fresh full MV-2 media. The following siRNAs were used: ON-TARGETplus SMARCA4/BRG1 siRNA (L-010431-00), ARID1A siRNA (L-017263-00) and Nontargeting pool siRNA (D-001810-10) from Dharmacon.

Transfection of HeLa cells was completed using a combination of OptiMEM Serum Reduced (Gibco), FuGENE HD Transfection Reagent (Promega), and plasmid DNA constructs for SOX18 wt, mutants or mChery. A mixture of OptiMEM and FuGENE HD (Promega) 4 µL/1000 ng DNA was created to a total volume of 100 µL, after which an appropriate amount of plasmid DNA was added. The mixture was then vortexed at 1000 rpm for a few seconds before being incubated at RT for 20 minutes. Then, the mixture was added to the cells with 900µl of fresh DMEM media per well and incubated at 37°C for total of 72 hours.

### BioID coupled with mass spectrometry

Protein-protein interaction screen BioID (Roux et al., 2018) was performed from stably KSHV-infected iSLK.219 and parental uninfected SLK cell line, transduced with BirA*SOX18 or BirA*mCherry construct containing lentiviruses. iSLK.219 was not induced, thus KSHV infection was strictly latent. Following transduction, cells were incubated and expanded for 72h before 80% full cultures were incubated with 50µM biotin (Pierce; B4639) for 24 hours. Cells were washed with PBS before scraping and pelleted before snap-frozen with liquid nitrogen and stored in −80°C. Next, the 1ml pellets were resuspended in 3x volume (3ml) of ice cold BioID lysis buffer (wash buffer (see below) with 0.1% SDS) and 1:3000 benzonase nuclease was added. The samples were vortexed and kept on ice for 15 min and sonicated with low output settings 45s on ice for 3 cycles, 5 min on ice in between. After, samples were centrifuged at 16.000g for 15 min at 4°C, and supernatants were transferred to new tubes and spin was repeated. From final supernatant, 50 µl inputs were removed for Western blot to check the expression of transduced plasmids. The supernatant samples were transferred through Bio-Spin chromatography columns (BioRad; 7326008), containing 200µl of Strep-Tactin Sepharose beads (IBA; 2-1201-002, 50% suspension), prewashed 3x 1ml with wash buffer (HENN-buffer with 0.5% IGEPAL, 1mM DTT, 1mM PMSF, 1,5mM Na_3_VO_4_, protease inhibitors; Sigma) for affinity purification. After supernatants were drained under gravity flow, the columns were washed four times with HENN-buffer (50mM HEPES pH 8.0, 5mM EDTA, 150mM NaCl, 50mM NaF, stored at 4°C in the dark). Then, the columns were closed, and biotin-bound proteins were eluted from the beads in the column with 300µl of fresh Biotin-HENN buffer (HENN-buffer with 0.5mM biotin) by incubating 5 min before opening the columns and flow-through was collected, and these steps were repeated. Final elution (600µl) was then frozen in −80°C before further processed and analyzed in Proteomics Unit (Institute of Biotechnology, University of Helsinki). Briefly, reduction of the cysteine bonds with 5mM Tris(2-carboxyethyl) phosphine (TCEP) for 30 mins at 37°C and alkylation with 10mM iodoacetamide was performed. The proteins were then digested to peptides with sequencing grade modified trypsin (Promega, #V5113) at 37°C overnight. After quenching with 10% TFA, the samples were desalted by C18 reversed-phase spin columns according to the manufacturer’s instructions (Harvard Apparatus). The eluted peptide sample was dried in vacuum centrifuge and reconstituted to a final volume of 30μl in 0.1% TFA and 1% CH_3_CN. BioID was performed with liquid chromatography-mass spectrometry (LC-MS) and analyzed as described in (Liu et al., 2018) by Proteomics Unit (Institute of Biotechnology, University of Helsinki). The high-confidence interacting proteins were identified by first filtering the data using Contaminant Repository for Affinity Purification (CRAPome) and Significance Analysis of INTeractome (SAINT)-express version 3.6.0. Then, only interacting proteins with ≥2 found unique peptides were selected, and finally BirA*SOX18 interacting proteins were bait-normalized to BirA*Cherry interacting proteins using the PSM (peptide spectral match) values.

### Cell viability assay

For measuring the viability of cells, CellTiter-Glo (Promega; G7572) luminescent viability assay (CTG) was performed on black 96-well ViewPlates (Revvity; 6005182) for 10 min and the luminescence from live cells were measured with FLUOstar Omega microplate reader (BMG Labtech). The viability % was calculated as an average of luminescent signal from triplicates comparing to DMSO treated cells considered as control with 100% viability.

### Cell proliferation assay

To compare the proliferation rates, low number of cells were plated on 96-well ViewPlates (Revvity; 6005182) and the next day the cells were treated with 10µM 5-ethynyl-2’-deoxyuridine EdU (Thermo Fisher) for 4 h with LEC/KLEC, and 2 h with HeLa cells, and fixed in 4% PFA in PBS. The proliferating cells were visualized using Click-iT EdU Alexa Fluor 647 (Molecular Probes; C10340) staining according to manufacturer’s instructions, and Hoechst 33342. Images were taken using Phenix Opera 20x and the portion of EdU-containing nuclei was quantified with Harmony software.

### Quantitative RT-qPCR

Total RNA was isolated from cells using the NucleoSpin RNA extraction kit (Macherey-Nagel; 740955) according to manufacturer’s protocols. Real time quantitative Polymerase chain reaction (RT-qPCR) and LightCycler480 PCR 384 multiwell plates (4titude FrameStar; 4ti-0382) were used to measure relative mRNA expression of samples. Primer sequences used to amplify the indicated targets are listed in Table 2. Relative abundances of viral mRNA were normalized by the delta threshold cycle method to the abundance of actin.

**Table 1.**
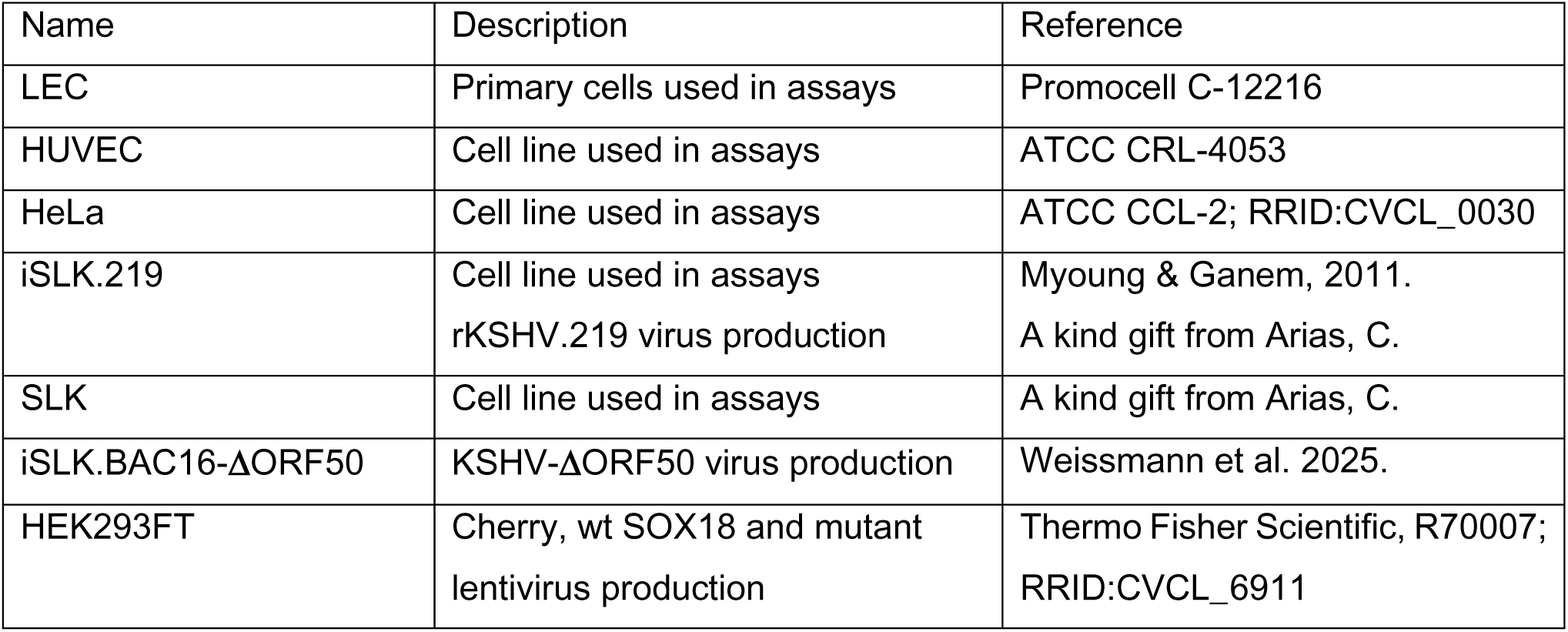
Cells used in this study.

| Name | Description | Reference |
| --- | --- | --- |
| LEC | Primary cells used in assays | Promocell C-12216 |
| HUVEC | Cell line used in assays | ATCC CRL-4053 |
| HeLa | Cell line used in assays | ATCC CCL-2; RRID:CVCL_0030 |
| iSLK.219 | Cell line used in assays<br>rKSHV.219 virus production | Myoung & Ganem, 2011.<br>A kind gift from Arias, C. |
| SLK | Cell line used in assays | A kind gift from Arias, C. |
| iSLK.BAC16-ΔORF50 | KSHV-ΔORF50 virus production | Weissmann et al. 2025. |
| HEK293FT | Cherry, wt SOX18 and mutant<br>lentivirus production | Thermo Fisher Scientific, R70007;<br>RRID:CVCL_6911 |

**Table 2.**
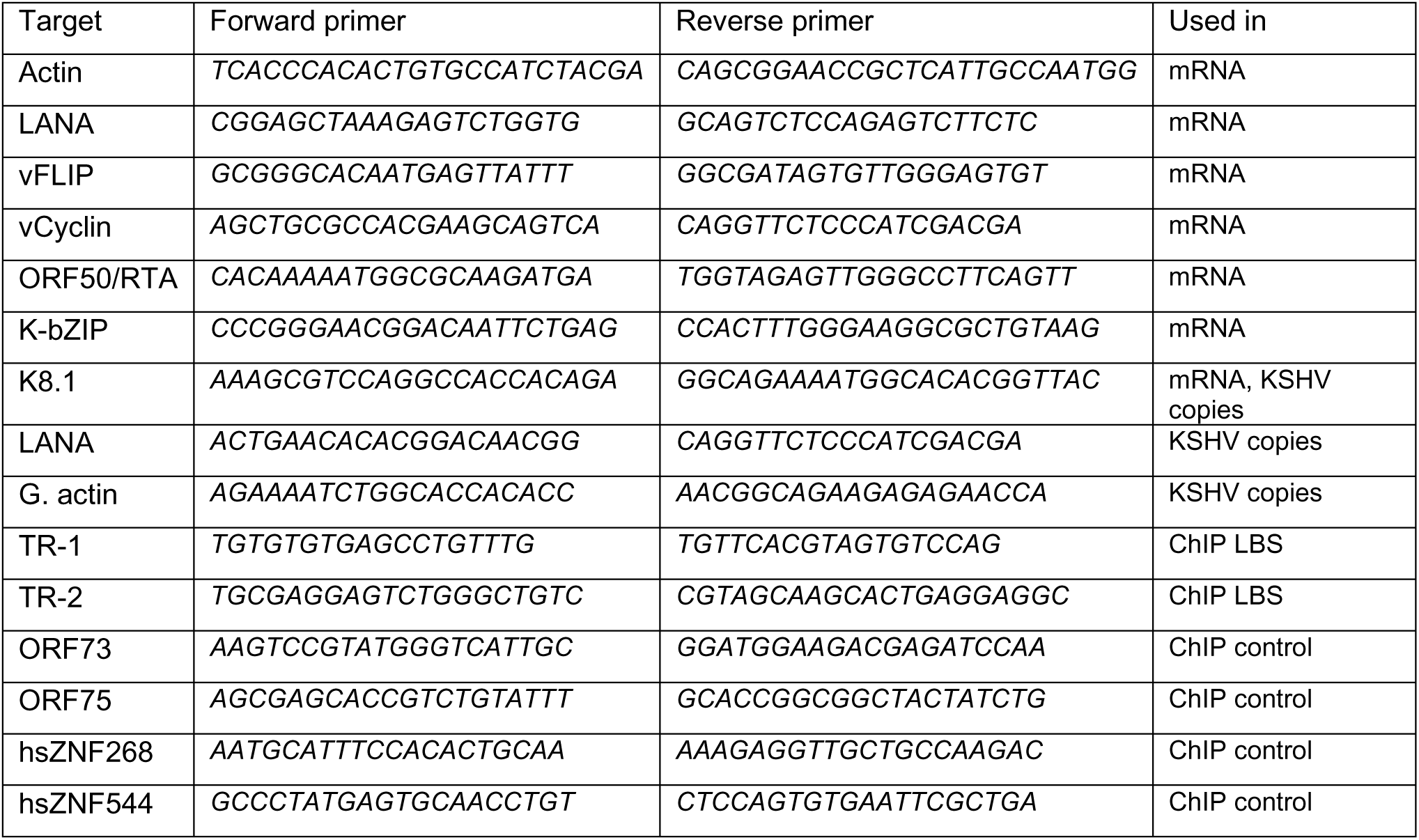
Oligonucleotides used in this study.

### Quantification of intracellular viral episome copies

Total DNA was isolated from cells using NucleoSpin Tissue Kit (Macherey-Nagel; 74098) and the KSHV genome episome copies were quantified by qPCR using 2XSYBR reaction mix (Fermentas; K0223) and unlabelled primers specific for LANA, K8.1, and genomic actin, listed in Table 2.

### Immunoblotting

Cell lysis, SDS-PGE and immunoblot were performed as described in (Gramolelli et al., 2018). The following primary antibodies were used for KSHV: rat monoclonal anti-HHV-8 LANA (Abcam; LN-35; ab4103); rabbit polyclonal RTA/ORF50 (a kind gift from C. Arias, University of CA), mouse monoclonal anti-K-bZIP and K8.1 (Santa Cruz; sc-69797, sc-65446) and for human: mouse monoclonal anti-β actin (Santa Cruz; sc-8432); mouse monoclonal anti-Vinculin (Santa Cruz; sc-73614); mouse monoclonal anti-SOX18 (Santa Cruz; sc-166025); rabbit monoclonal anti-ARID1A and anti-BRG1 (Abcam; ab182560, EPR13501 and ab110641, EPNCIR111A); anti-H2A (Cell Signaling; D6O3A, 12349S) or anti-H2B (Cell Signaling; D2H6, 12364S) and mouse monoclonal anti-HP1α (Santa Cruz; sc-515341). Following secondary antibodies were used: anti-mouse, anti-rabbit and anti-rat IgG HRP conjugated (Cell Signaling; 7076, 7074, 7077).

### Proximity ligation assay (PLA)

Proximity ligation assay (PLA) was performed using Duolink PLA technology (Sigma-Aldrich). LECs and KLECs were plated on a PhenoPlate (Revvity; 6055300) and infected with rKSHV.219. Cells were fixed with 4% PFA then permeabilized with Triton X-100 (Sigma; T9284) and 1µg/mL of Hoechst 33342 (Fluka Biochemicka) in PBS. Blocking with Duolink Blocking Solution in a 37°C humidity chamber for 60 minutes and then stained with antibodies against rabbit polyclonal anti-LANA (a kind gift from B. Chandran lab), mouse monoclonal anti-SOX18 (Santa Cruz; sc-166025), anti-BRG1 (sc-17796), or anti-ARID1A (sc-32761) or either normal mouse IgG or normal rabbit IgG (sc-2025; sc2027). Wells were washed five times with 1x wash buffer A and then treated with PLA probe solution composed of anti-mouse PLUS (DUO92001) and anti-rabbit MINUS (DUO92005) probes diluted in Duolink antibody diluent and incubated in a 37°C humidity chamber for 60 minutes. Probes were detected with *in situ* far-red detection reagent (DUO92013). Ligation was performed by treating cells with 1:40 dilution of Ligase in 1x Ligation Buffer and incubating and 37°C humidity chamber for 30 minutes. Wells were washed five times with 1x wash buffer A and Amplification was performed by treating cells with 1:80 dilution of Ligase in 1x ligation buffer and incubating and 37°C humidity chamber for 100 minutes. Wells were washed five times with 1x wash buffer B and a 0.01x wash buffer B was added. Imaging of interaction PLA dot signals were accomplished using Opera Phenix (PerkinElmer) and quantified using Harmony software.

### Chromatin immunoprecipitation (ChIP)

HeLa cells transfected with Cherry, SOX18, C240X, or HMGdel cDNAs, or LECs were infected with rKSHV.219. After 72 hours, the infection efficiency was confirmed by GFP signal. For KLEC, cells were treated with inhibitors or DMSO control and incubated for another 24 hours. For each ChIP, one or three 10-cm dish for HeLa and KLEC, respectively, was cross-linked and protocol according to SimpleChIP Enzymatic Chromatin IP Kit (Cell Signaling; 9003S) was used. Antibodies against rat monoclonal anti-HHV-8 LANA (Abcam; LN-35, ab4103) and IgG control (Cell Signaling; 9003S) and were used, also listed in Table 3. Chromatin was eluted and de-crosslinked and DNA was purified using a DNA Clean & Concentrator kit (Zymo Research; D5205). The experiments were done at least two independent times. The purified DNA was amplified with RT-qPCR with two set of TR primers (mean) on known LBS (LANA-binding sites), and primers for non-LBS on KSHV genome as well as two additional primers for human genome as negative controls listed in Table 2, and differences in samples is shown as % of input as individual values for each biological replicate.

**Table 3.** Antibodies used in this study.

| Antibody stain | Species | Source/reference | Dilution | Used in |
| --- | --- | --- | --- | --- |
| BrdU | Mouse monoclonal | BD Biosciences, 555627 | 1:60 | BrdU-IP |
| LANA | Rat monoclonal | Abcam, ab4103 | 1:1000<br>1:500 | IF<br>WB |
| LANA | Rabbit | A kind gift from B. Chandran<br>University of | 1:1000 | PLA |
| ORF50/RTA | Rabbit polyclonal | A kind gift from C. Arias,<br>University of CA | 1:1000 | WB |
| K-bZIP | Mouse monoclonal | Santa Cruz, sc-69797 | 1:1000 | WB |
| K8.1 | Mouse monoclonal | Santa Cruz, sc-65446 | 1:1000 | WB |
| β-actin | Mouse monoclonal | Santa Cruz, sc-47778 | 1:1000 | WB |
| Vinculin | Mouse monoclonal | Santa Cruz, sc-73614 | 1:1000 | WB |
| Mouse IgG | Mouse | Santa Cruz, sc-2025 | 1:250<br>1:1000 | BrdU-IP<br>PLA |
| Rabbit IgG | Rabbit | Cell Signaling 2729S | 1:1000 | PLA |
| BrdU | Mouse monoclonal | BD Biosciences, 555627 | 1:60 | BrdU-IP |
| SOX18 | Mouse monoclonal | Santa Cruz, sc-166025 | 1:1000 | IF, WB, PLA |
| ARID1A | Rabbit monoclonal | Abcam, ab182560 | 1:500 | IF, WB, PLA |
| ARID1A | Mouse monoclonal | Santa Cruz, sc-32761 | 1:1000 | PLA |
| BRG1 | Rabbit monoclonal | Abcam, ab110641 | 1:500 | IF, WB, PLA |
| BRG1 | Mouse monoclonal | Santa Cruz, sc-17796 | 1:1000 | PLA |
| HP1α | Mouse monoclonal | Santa Cruz, sc-515341 | 1:1000 | IF |
| H2A | Rabbit monoclonal | Cell Signaling, 12349S | 1:1000 | IF, WB |
| H2B | Rabbit monoclonal | Cell Signaling, 12364S | 1:1000 | WB |

### Bromodeoxyuridine (BrdU) incorporation assay

HeLa cells transfected with Cherry, SOX18, C240X, or HMGdel cDNAs, or LECs were infected with rKSHV.219. After 72 hours, the infection efficiency was confirmed by GFP signal. For KLEC, cells were treated and incubated for another 72 hours. Cell culture media with 100µM of BrdU (Sigma; B-5002) was added and incubated for 2 hours for HeLa and 4 hours for KLECs before cell samples were trypsinized and collected. The samples were centrifuged for 5 minutes at 1500 rpm to collect a cell pellet. The cell pellet was then used for DNA extraction using the Nucleospin Tissue kit (Macherey-Nagel; 740952.250) with T1 lysis buffer and Proteinase K. DNA samples were eluted and sonicated for 3 cycles for 30 seconds, with the samples cooling on ice between each cycle. For BrdU pulldown, DNA was first denatured at 95°C for 10 minutes and cooled on ice. For input samples, 10% of each sample was aliquoted and stored. Following, 4µL of mouse monoclonal BrdU antibody (BD Biosciences; 555627) or normal mouse IgG (Santa Cruz; sc-2025) was added to the remaining samples and incubated overnight at 4°C while rotating. Additionally, a total of 130µL of DynabeadsTM Protein G magnetic beads (Invitrogen; 10003D) were washed twice with 1mL IP wash buffer (50mM HEPES-NaOH pH 7.55, 250mM LiCl, 1mM EDTA, 1% NP-40, 0.7% sodium deoxycholate) and once with IP buffer (10mM HEPES-NaOH pH 7.9, 100mM NaCl, 1mM EDTA, 0.5 mM EGTA, 0.1% sodium deoxycholate) with the magnetic rack (Bio-Rad). The beads were then blocked with 1mg/mL of BSA and 0.25mg/mL of Salmon sperm DNA (Fisher; 10605543) overnight at 4°C, rotating. The next day, the blocking buffer in the magnetic beads were removed using the magnetic rack and washed once with IP buffer. Then, 130µL of IP buffer were mixed by pipetting and 15µL of the beads were added to each sample and mixed. The samples were then incubated for 3 hours at RT, rotating. Following, the beads in the samples were washed 5 times with IP wash buffer using 1mL for each sample. The samples were then eluted in 200µL of elution buffer (50mM Tris-HCl pH 8, 10mM EDTA, 1% SDS) at 65°C for 1 hour. Following elution, the samples were added to the magnetic stand and the supernatant was transferred to new, clean tubes. The samples and inputs were then purified with the ChIP DNA Clean and Concentrator kit (Zymo Research; D5205). A total of 40µL of elution buffer was used for the final DNA product for a subsequent RT-qPCR run conducted using primers for KSHV and the human housekeeping genes of interest listed in Table 2 and the differences in samples are shown as % of BrdU incorporation normalized to human BrdU.

### Antibodies

Primary antibodies used in Western Blotting (WB), immunofluorescence staining (IF), Proximity Ligation Assay (PLA), chromatin immunoprecipitation (ChIP), and bromodeoxyuridine immunoprecipitation (BrdU-IP) assays of this study are listed in Table 3.

### Image acquisition for LECs

#### Immunofluorescence

Cells were plated on a 96-well PhenoPlate (Revvity; 6055300) and infected with rKSHV.219 and treated or cells were seeded on fibronectin (from human plasma, Sigma; F0895) -coated glass coverslips on 24-well plate, and treated. Cells were fixed with 4% PFA, permeabilized with Triton X-100 (Sigma; T9284) and stained for KSHV proteins with antibody against rat monoclonal anti-HHV-8 LANA (Abcam; LN-35; ab4103) and for human proteins with antibodies against mouse monoclonal anti-SOX18 (Santa Cruz; sc-166025), rabbit monoclonal anti-BRG1 (Abcam; EPNCIR111A) or anti-ARID1A (EPR13501), anti-H2A (Cell Signaling; D6O3A, 12349S) or mouse monoclonal anti-HP1α (Santa Cruz; sc-515341). Alexa Fluor anti-rat 647 or anti-mouse 596 (Invitrogen; A48272, A21203) were used as secondary antibodies. Nuclei were visualized with Hoechst 33342 (Sigma; 14533).

#### High-throughput imaging

From 96-well plates, images were taken using an automated cell imaging system ImageXpress Pico (Molecular Devices) with 10x objective or Opera Phenix (PerkinElmer) with either 20x or 40x objectives with z-stack of 5 images. Cells were quantified using pipeline created in Harmony. Briefly, LANA speckles were quantified as mean number of nuclear objects from 10 fields (n > 200 nuclei) for each biological replicate (n = 3) shown as individual values ±SD. Signal from PLA dots were quantified as mean number of nuclear objects from 10 fields (n = 100 nuclei) combined from (n=3) biological replicates shown as violin plots with median and quantiles.

#### Confocal

Coverslip images were taken with LSM880 (Zeiss) PMT confocal with Plan-Apochromat 63x oil objective with 405, 561 and 633nm lasers. Cells were quantified using macro pipeline created in Fiji-imageJ. Briefly, maximum image projections (MIP) were created from z-stack of 15 images for each channel. Signal intensity thresholds were acquired for 16-bit depth MIP images with default settings, and particles (nuclei) were analyzed for mean arbitrary unit (a.u.) intensity within each nucleus (n = 100 or 200) shown as individual values ±SD. LANA speckles were quantified as mean number of nuclear objects in each nucleus (n = 100) shown as violin plots with median and quantiles.

### Image acquisition for HUVEC

#### Confocal

HUVEC cells were seeded on 0.5% gelatin coated 8-well microscope slide (Ibidi; 80827) at a density of 50,000 cells/well in EBM-2 media overnight. HUVEC cells were then treated overnight with either DMSO or Sm4 (30µM). The next day cells were stained with SiR-DNA (Spirochrome) at a 1:2000 dilution for 1 hour prior to imaging. Cells were imaged on a Leica TCS SP8 (Leica Microsystems GmbH) microscope at 37°C and 5% CO_2_ using a 93x 1.30NA glycerol immersion objective and a tunable white light laser unit.

#### STED

Live cell STED gated imaging was performed on a Leica TCS SP8 microscope (Leica Microsystems), using a 93x 1.30 NA glycerol immersion motCORR STED White objective and a tunable white light laser unit. Spy650-DNA labelled nuclei were imaged with a 647nm excitation running at ∼5% power, and with emission wavelengths set between 660 and 750nm. Fluorescence depletion used a 775nm laser running at 50% output power. Pinhole was set at 129.5μm. Emitted fluorescence intensity was filtered by a notch filter (775nm). Images (1024 by 1024 pixels) were collected with a pixel size range of ∼ 25nm – 30nm (Zoom range between 4 to 5) and with a Z-stack step size of 0.15μm (typical range of 0.3 – 0.5μm). A line average of 3 with no frame accumulation was used. All scans were performed at a scan speed of 400 Hz. Deconvolution of images was performed with Huygens Professional software (Scientific Volume Imaging). Average intensity of the samples was analyzed on cell profiler and the line profiles were determined using ImageJ.

### Microscopic imaging of epigenetic landscape MIEL

#### Preparation

HUVEC cells were seeded at a density of 50,000 cells/well in an 8-well chamber slide (Ibidi; 80827). Cells were then treated with either DMSO or Sm4 overnight. LECs were seeded at a density of 100,000 cells/well in a 12-well plate and the next day either left uninfected or infected with rKSHV.219. The next day cells were moved to fibronectin (from human plasma, Sigma; F0895) coated glass coverslips on 24-well plate. After 72 hours post infection, cells were then treated with either DMSO or Sm4 for 24 hours.

The following day both ECs were fixed with 4% PFA for 10 mins at room temperature, then washed with PBS. Cells were then permeabilized with 0.3% Triton X-100/PBS for 5 minutes, followed by washing with PBS and blocking with 0.5% BSA/PBS for 1 hour. Cells were then stained with 5 ug/ml of DAPI for 3 minutes and then washed with PBS. LECs and KLECs were additionally stained with mouse monoclonal anti-HP1α (Santa Cruz; sc-515341) overnight, washed with PBS and Alexa Fluor anti-mouse 596 (Invitrogen; A21203) was used as secondary antibody. Coverslips were washed with PBS and dH2O before mounting to microscope slides.

#### Confocal acquisition

HUVEC cells were acquired on a Nikon A1R confocal microscope using the 20x 0.75 NA air objective and LECs/KLECs were acquired on a Zeiss LSM880 confocal microscope using the 63x oil objective.

#### Data Processing

Image features for each cell were extracted using the MIEL pipeline (Farhy et al., 2019). Feature values were normalized using z-score transformation. For each experimental condition, individual cell profiles were condensed into an averaged center representing the population-level feature vector. The number of cells used to compute each averaged center was determined through bootstrap analysis, as described in Bootstrap analysis. The condensed centers were then subjected to principal component analysis (PCA), using four principal components, to construct a reduced-dimensional representation of cellular behavior.

#### Distance Matrix

To assess phenotypic similarity between conditions, the Euclidean distance between all pairwise centers was calculated. To evaluate within-condition variability, the average distance between all centers belonging to the same experimental condition was computed, providing a measure of dispersion in cellular behavior across replicates.

#### Confusion Matrix

Support Vector Machine (SVM) classification was performed using Python’s scikit-learn library (version 1.2.2) to evaluate the separability of the condensed centers. Eighty percent of the data from each condition were used as a training set, while the remaining 20% served as the test set. Classification accuracy was assessed on the test set, and results were summarized in a confusion matrix to visualize performance across conditions.

#### Bootstrap Analysis

To determine the optimal number of cells required to generate a representative averaged center, we conducted a bootstrap-based optimization. First, PCA was performed using all available cells from all conditions. From this analysis, we selected conditions that exhibited clear separation in the PCA space. Using these separable conditions, 1,000 bootstrap iterations were performed. In each iteration, 80% of the total cell count per condition was randomly sampled with replacement. These subsampled datasets were condensed into centers, and PCA followed by SVM classification was applied. Classification accuracy was recorded for each iteration. The optimal condensation number was defined as the smallest number of cells that achieved ≥95% classification accuracy in ≥95% of the bootstrap iterations. This ensured robust and reproducible discrimination of experimental conditions while minimizing cell input requirements.

### Quantitative Molecular Imaging

#### Single molecule tracking (SMT)

##### Preparation

SMT was performed as described in (McCann et al., 2021). In summary, HeLa cells were seeded at a density of 23,000 cells/well in 8-well chamber glass slides (Ibidi; 80827) coated with 0.5% gelatine 24h prior to transfection. 300ng of plasmid DNA/well of either Halo-tagged SOX18 was transiently transfected into the cells using X-tremeGENE 9 Transfection Reagent kit (Roche; XTG9RO). Cells were then incubated at 37°C with 5% CO_2_ overnight prior to imaging then washed three times. Culture media was replaced with imaging media (FluoroBrite DMEM, Gibco; A1896701), 10% FBS, 1% HEPES, 1% GlutaMax along with 400nM of Trichostatin A for four hours prior to imaging. 45 minutes before imaging 1nM of JF549 Halo-tag ligand (Promega; GA1111) was added directly to the media and cells were incubated for 10 minutes at 37 ° C with 5% of CO_2_. Following incubation, cells were washed twice 15 minutes apart and replaced with imaging media.

##### SMT acquisition

Images were acquired on a Nikon TIRF microscope at a TIRF angle of 61 degrees to achieve HILO illumination. Samples were recorded with an iXon Ultra 888 EMCCD camera, filter cube TRF49909 – ET – 561 laser bandpass filter and 100 X oil 1.49 NA TIRF objective. Cells were imaged using a 561nm excitation laser at a power density of 10.3μW to perform two different acquisition techniques. A fast frame rate which uses a 50 Hz (20ms acquisition speed) to acquire 6000 frames without intervals to measure displacement distribution and fraction bound, and a slow frame rate which uses a 2 Hz (500ms acquisition speed) to acquire 500 frames without intervals to measure residence times.

##### SMT analysis

Masking and segmentation of the nucleus was performed in ImageJ for all files. To identify and track molecules a custom-written MATLAB implementation of the multiple target tracing (MTT) algorithm, known as SLIMfast was used (Serge et al., 2008). Parameters used for fast frame rate analysis: Localization error: 10^-6.5^, blinking (frames) = 1, max # of competitors: 3, max expected diffusion coefficient = 3μm^2^/sec, box size = 7, timepoints = 7, clip factor = 4. Cells with less than 500 trajectories based on the above parameters were excluded from analysis. The first four frames of each trajectory were used to calculate the mean squared displacement. Diffusion coefficient was calculated from each trajectory’s mean squared displacement and plotted. An inflection point was determined on SOX18 diffusion profile and used as a boundary to determine confined vs non-confined states for all conditions. The confined fraction and non-confined fraction of each cell were calculated by computing the number of trajectories whose mean diffusion coefficient were defined as confined and non-confined respectively. Parameters used for slow frame rate analysis: Localization error: 10^-7^, blinking (frames): 1, max # of competitors: 3, max expected diffusion coefficient = 0.33μm^2^/sec, box size = 9. Slow-tracking analysis was performed using custom MATLAB code based on (Chen et al., 2014).

### Fluorescence fluctuation spectroscopy (FFS)

#### Halo Tag labelling for FFS experiment

To saturate all Halo Tags with Janelia Farm dyes 549 or 646 for single or dual-color labelling, HeLa cells transiently expressing Halo Tag-tagged Sox7 and SOX18 constructs were incubated with a saturating concentration of a single JF dye (JF549, 100nM) or two JF dyes (JF549 : JF646, 100nM : 100nM) for 15 minutes at 37°C, similar to (McCann et al., 2021). Cells were washed twice with 1× PBS before imaging experiments.

#### FFS acquisition

All FFS measurements for Number and Brightness (NB) analysis and cross Raster Image Correlation Spectroscopy (RICS) were performed on an Olympus FV3000 laser scanning microscope coupled to an ISS A320 Fast FLIM box for fluorescence fluctuation data acquisition. For single channel NB FFS measurements, HaloJF549 tagged plasmids were excited by a solid-state laser diode operating at 561 nm and the resulting fluorescence signal was directed through a 405/488/561 dichroic mirror to an external photomultiplier detector (H7422P-40 of Hamamatsu) fitted with a mCherry 600-640 nm bandwidth filter. For dual channel RICS FFS measurements (that enable cross RICS), the HaloJF546 and HaloJF646 combination were excited by solid-state laser diodes operating at 561 nm and 640 nm, respectively, and the resulting signal was directed through a 405/488/561/640 dichroic mirror to two internal GaAsP photomultiplier detectors set to collect 600-640 nm and 650-750 nm, respectively.

All FFS data acquisitions (i.e., NB, cross RICS) employed a 60X water immersion objective (1.2 NA) and first involved selecting a 10.6 μm region of interest (ROI) within a HeLa cell nucleus at 37°C in 5% CO2 that exhibited low protein expression level (nanomolar) to ensure observation of fluctuations in fluorescence intensity. Then a single or simultaneous two channel frame scan acquisition was acquired (N = 100 frames) in the selected ROI with a pixel frame size of 256 x 256 (i.e., pixel size ∼ 41nm) and a pixel dwell time of 12.5 µs. These conditions resulted in scanning pattern that was found to be optimal for simultaneous capture of the apparent brightness and mobility of the Halo tagged constructs being characterised by NB and cross RICS analysis; all of which was performed in the SimFCS software developed at the Laboratory for Fluorescence Dynamics (LFD).

#### Number and brightness (NB) analysis

The oligomeric state of the different HaloJF549-tagged plasmids investigated was extracted and spatially mapped throughout single channel FFS measurements via a moment-based brightness analysis that has been described in previously published papers ^79,80^. In brief, within each pixel of an

NB FFS measurement there is an intensity fluctuation F(t) which has: (1) an average intensity 〈F(t)〉 (first moment) and (2) variance 𝜎^2^ (second moment); and the ratio of these two properties describes the apparent brightness (B) of the molecules that give rise to the intensity fluctuation. The true molecular brightness (*ɛ*) of the fluorescent molecules being measured is related to B by B = ε + 1, where 1 is the brightness contribution of a photon counting detector. Thus, if we measure the B of monomeric HaloJF549-Sox7 (B_monomer_= *ɛ_monomer_* + 1) under our NB FFS measurement conditions, then we can determine *ɛ_monomer_* and extrapolate the expected B of HaloJF549-tagged dimers (B_dimer_= *(2 x ɛ_monomer_)* + 1) or oligomers (e.g., B_tetramer_= *(4 x ɛ_monomer_) + 1),* and in turn define brightness cursors, to extract and spatially map the fraction of pixels within a NB FFS measurement that contain these different species. These cursors were used to extract the fraction of HaloJF549-SOX18 dimer and oligomer (i.e., number of pixels assigned B_dimer_ or B_oligomer_) within a NB FFS measurement and quantify the degree of SOX18 self-association across multiple cells. Artefact due to cell movement or photobleaching were subtracted from acquired intensity fluctuations via use of a moving average algorithm and all brightness analysis was carried out in SimFCS from the Laboratory for Fluorescence Dynamics.

### Assay for Transposase-Accessible Chromatin -sequencing (ATAC-seq)

#### Cell preparation and infection comparison of KSHV strains

LECs infected with rKSHV.219 display a unique infection program with spontaneous lytic reactivation initiated by viral ORF50/RTA, leading to production of viral progeny (Choi et al., 2020; Gramolelli et al., 2020). These viral particles contain nascent DNA that, upon lysis of the host cell, releases cell free viral DNA that interferes with the ATAC-seq analysis resulting in high background signal. To avoid this, we opted to infect LECs with a KSHV-BAC16-ΔORF50 strain that has ORF50/RTA stop-codon prohibiting spontaneous lytic reactivation, and production of new viral particles (Weissmann et al., 2025). Notably, although KSHV-ΔORF50 is an optimal viral strain for ATAC-seq, most of the infection assays in this study are carried out using rKSHV.219 strain as it better recapitulates the natural infection of KSHV in LECs.

Infection by ΔORF50 was first validated to induce similarly high levels of intranuclear episome copies and hallmarks of KSHV infection in LECs as wt rKSHV.219 (Fig S4A-F). Importantly, the SOX18 upregulation is also evident in ΔORF50-KLECs (Fig S4A). CTG viability assay using increasing concentrations of Sm4 to compare the viability of LECs infected with both strains show that infection sensitizes LECs to Sm4 and that 20µM Sm4 is optimal for all SOX18 inhibition analyzes in LECs, including ATAC-seq (Fig S4B). Infections in LECs resulted in identical spindling phenotypes, a hallmark of KSHV-infected LECs, which were similarly reduced upon SOX18 blockade by Sm4 (Fig S4C). Next, we checked the relative intracellular KSHV episome numbers by qPCR, which were reduced by Sm4 to the same extent (Fig S4D). LANA protein tethers KSHV episomes to host chromatin and its characteristic dotty IF staining pattern (LANA speckles) can therefore serve as a surrogate marker for intranuclear viral episomes (Adang et al., 2006). LECs infected with either viral strain showed comparable levels of nuclear LANA speckles (Fig S4E, left panels) confirming that infections yield similar levels of intranuclear episome copies in LECs. The amount of LANA speckles was similarly reduced by Sm4 in LECs (Fig S4E-F).

#### Library preparation for sequencing

ATAC-seq libraries for uninfected LEC and ΔORF50-KLEC treated for 24h with DMSO or Sm4 were prepared in B. Sahu lab as previously described in (Buenrostro et al., 2015; Corces et al., 2017). Briefly, 50,000 cryopreserved cells were washed with ice-cold PBS and resuspended in 50µl of ATAC-seq lysis buffer and incubated for 3 min on ice. Nuclei were centrifuged at 500 x g for 10 min at 4°C, followed by transposition with Tn5 transposase (Illumina; 20034197). Tagmentation was carried out on a thermomixer at 37°C for 30 min at 1,000 rpm. The reaction was purified using MinElute PCR Purification Kit (Qiagen; 28004) and eluted in nuclease-free water. The samples were amplified for a total of 8 cycles and purified with AMPure beads (Agencourt; A63881). Libraries were paired end sequenced on Illumina Novaseq 6000.

ATAC-seq for ΔORF50-KLEC treated for 72h with DMSO, Sm4 or FHT-1015, was performed in A. Grunhoff lab using the Omni-ATAC-seq protocol (Corces et al., 2017). Briefly, 50,000 cryopreserved cells were thawed, treated with DNase I (200U/ml, Worthington) at 4°C for 5 min and DNAse was inactivated by addition of EDTA (1.5mM final). Cells were washed with cold wash buffer (PBS + 0.04% BSA) twice and 1×10^5^ cells were resuspended in 1ml cold RSB buffer (10mM Tris-HCl pH 7.4, 10mM NaCl, 3mM MgCl_2_). Cells were pelleted again at 500 x g for 5 min and resuspended in 50µl of cold ATAC-NTD lysis buffer (RSB Buffer + 0.1% NP40, 0.1% Tween20, 0.01% Digitonin). Lysed cells were diluted in 1ml cold ATAC-T buffer (RSB + 0.1% Tween20) and inverted three times. The resulting nuclei were pelleted at 500 x g for 10 minutes and the supernatant was removed. Cell pellets were transposed with 50µl of transposition mix containing 25µl 2xTD Buffer (20mM 1M Tris-HCl pH 7.6, 10mM MgCl_2_, 20% Dimethyl Formamide) 2.5µl transposase (custom made, 100nM final), 16.5µl PBS, 0.5µl 1% digitonin, 0.5µl 10% Tween-20 and 5µl H2O) at 37°C and 1000 rpm on a thermomixer for 30 min. The reaction was stopped by adding 250µl of DNA Binding Buffer and DNA was isolated using the Clean and Concentrator-5 Kit (Zymo; D4013). Libraries were produced by PCR amplification of tagmented DNA and sequenced on a NextSeq 2000 sequencer 50bp Paired End.

#### Bioinformatic analysis

The ATAC-seq data processed as previously described (Fei et al., 2023) and visualized with RPKM normalization and a binsize of 10. Briefly, for mapping of ATAC-seq data to both human genome and virus genome, we constructed a hybrid genome that included the hg38/GRCh38 version of human genome and Kaposi sarcoma herpesvirus genome (GenBank id: HQ404500.1) (referred as hybrid genome from now on). This hybrid genome included human chromosomes 1-21, X, Y and KSHV genome. The hybrid genome was used for all the ATAC-seq analysis steps. Briefly, Pearson correlation between the samples was calculated and visualized using deeptools (v.3.1.3) with outlier removal. Differential analysis of the chromatin accessibility in the ATAC-seq samples was done in R using Diffbind (v3.16.0). The analysis was conducted using alignment files and narrowPeak files. Sites with a false discovery rate (FDR) value of less than 0.05 were defined as differentially accessible. Differential site locations were compared using bedtools and visualized as heatmaps using deeptools with bigwig files. ATAC-seq signal in the viral genome was visualized using pyGenomeTracks (v3.9) with bigwig files. For the visualization of the human genome sites, the bigwig files were converted into bedGraph format using bigWigToBedGraph (v377) and the genomic coordinates were plotted using Spark (v2.6.2). Homer v4.10.4 was used to perform de novo motif analysis on the differential ATAC-seq sites. The chromatin accessibility loss and gain sites were ranked according to their log2FoldChange and up to 1000 differential accessibility sites with the highest fold change were selected for the motif analysis. findMotifsGenome.pl script was used to run de novo motif analysis with the hybrid genome using default parameters. Transcription factor Occupancy prediction By Investigation of ATAC-seq Signal (TOBIAS, v0.13.3) was used to predict transcription factor binding differences as previously described in (Fei et al., 2023). The gain and loss groups are defined as the TFs having -log10(p-value) above the 95% quantile or differential binding scores in the bottom or top 5% of the scores. Motifs were retrieved from Jaspar database. (https://jaspar.elixir.no/download/data/2024/CORE/JASPAR2024_CORE_vertebrates_non-redundant_pfms_jaspar.txt) and the results were visualized using ggplot2 (v3.5.1).

### Statistical analysis

Graphical presentations and statistical analysis were generated with GraphPad Prism Software v9.0 (Dotmatics). For statistical evaluation of the RT-qPCR data for relative KSHV genome copies, the logarithmic values were converted to linear log2 scale values by using the double delta CT (2-ΔΔ CT) method. Human genomic actin when measuring DNA, and actin when measuring mRNA were used as internal control and accounted in the calculations to correct differences in the RNA and DNA amount. The data is presented as individual values ± standard deviation (SD) between biological replicates unless otherwise reported. Statistical differences between groups were evaluated with either Student’s *t*-test (two-tailed) or Welch’s *t*-test, or ordinary one-way ANOVA followed by Dunnett or Tukey correction for multiple comparisons. Further details can be found from figure texts with p*-*values considered significant indicated by asterisk.

## Resource availability

Further information and requests for reagents may be directed to and will be fulfilled by the lead contacts PMO and MF.

## Materials availability

This study did not generate new unique reagents.

## Data and code availability

Datasets have been deposited at DOI: 10.5281/zenodo.15751062 and are publicly available as of the date of publication. The DOIs are listed in the key resources table.

---

**Supplementary Figure 1. Related to Fig 1.**
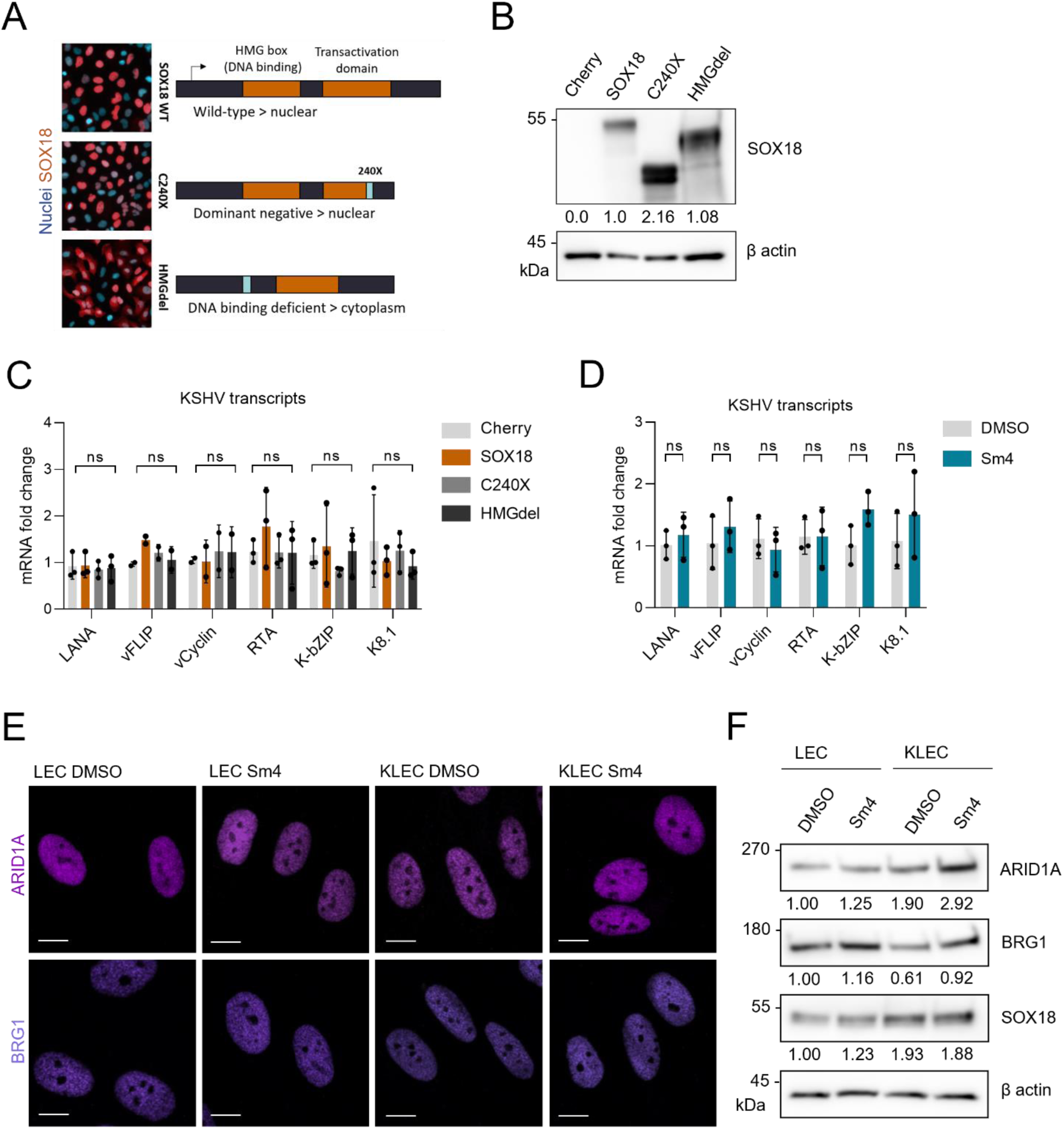
**A-C.** HeLa cells expressing SOX18wt, mutants C240X (dominant-negative transactivation deficient) or HMGdel (DNA-binding deficient), or mCherry as a control, and thereafter infected with rKSHV.219 for 72h (KSHV-HeLa). **A.** IF images of the SOX18wt and mutants expressing cells labeled with anti-SOX18 antibody and a schematic of the constructs. **B.** Immunoblotting with anti-SOX18 antibody using β-actin as a loading control for normalization. **C.** RT-qPCR for the indicated viral genes in KSHV-HeLa. **D.** LECs infected with rKSHV.219 (KLECs) for 72 hours and treated with Sm4 or DMSO control for 24h and relative mRNA measured for indicated viral transcripts. Statistical significance was determined by one-way ANOVA with Dunnett correction for multiple comparisons; ns = non-significant. **E-F.** Uninfected LECs and KLECs 72h p.i. treated with DMSO or Sm4 for another 72h and E) labeled with anti-ARID1A and -BRG1 antibodies, nuclei were counterstained with Hoechst (33342), scale bar is 10µm, and F) immunoblotted for the indicated proteins and quantified as in B.

**Supplementary Figure 2. Related to Fig 2.**
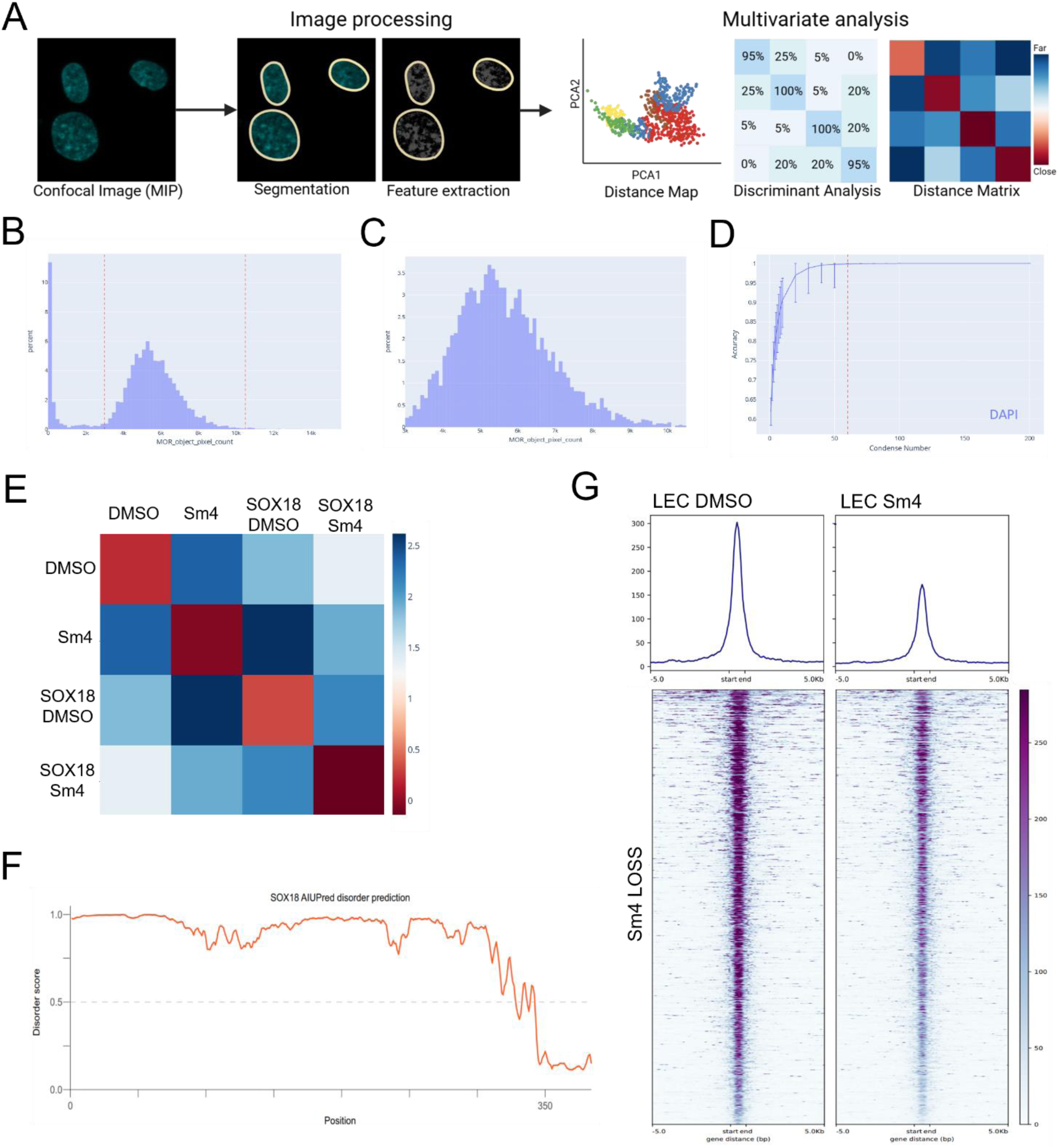
**A.** Representative pipeline of unsupervised MIEL analysis. For each experimental condition, confocal images of the total cell population are acquired and separated by fluorescence channel. Individual nuclei are segmented using DAPI staining as a mask. Within each segmented nucleus, 253 texture- and edge-based fluorescent features are extracted at the pixel level. These features are computed using the intensity relationships between each pixel and its surrounding neighbors, capturing spatial variation in signal distribution. The variation in the extracted nuclear features is quantified across all nuclei, and dimensionality reduction is performed using Principal Component Analysis (PCA). The top two principal components are used to visualize data structure and spread in a 2D PCA plot. To classify cell populations, a Support Vector Machine (SVM) algorithm is applied, enabling the identification of distinct clusters based on feature signatures. The average pairwise distances points in PCA space are then computed and represented as a similarity matrix. **B-C.** HUVECs ± SOX18 over-expression and ± Sm4 treatment stained with DAPI. **B.** Histogram displaying distribution of all object sizes (pixels) identified during segmentation, objects inside of red-dashed lines are used in analysis and **C.** zoom in on dashed lines in right panel. **D.** Line plots show accuracy measurements versus cell condense number; 95% confidence intervals are shown with red dotted line denotes smallest condense number above 95% accuracy. **E.** Average distance matrix calculated from the distance between each point per condition, with blue as farthest distances and red as closest distances. **F.** Intrinsic disorder prediction using AIUPred for SOX18 transcription factor (Uniprot ID: P35713). Y-axis represents the disorder prediction score and x-axis amino acid position. Scores over 0.5 (dashed line) are considered disordered. **G.** LECs treated with DMSO or Sm4 and subjected to ATAC-seq. Heatmap showing chromatin accessibility loss on the top 1000 sites of host genome (dark blue maps and line) upon Sm4 treatment.

**Supplementary Figure 3. Related to Fig 3.**
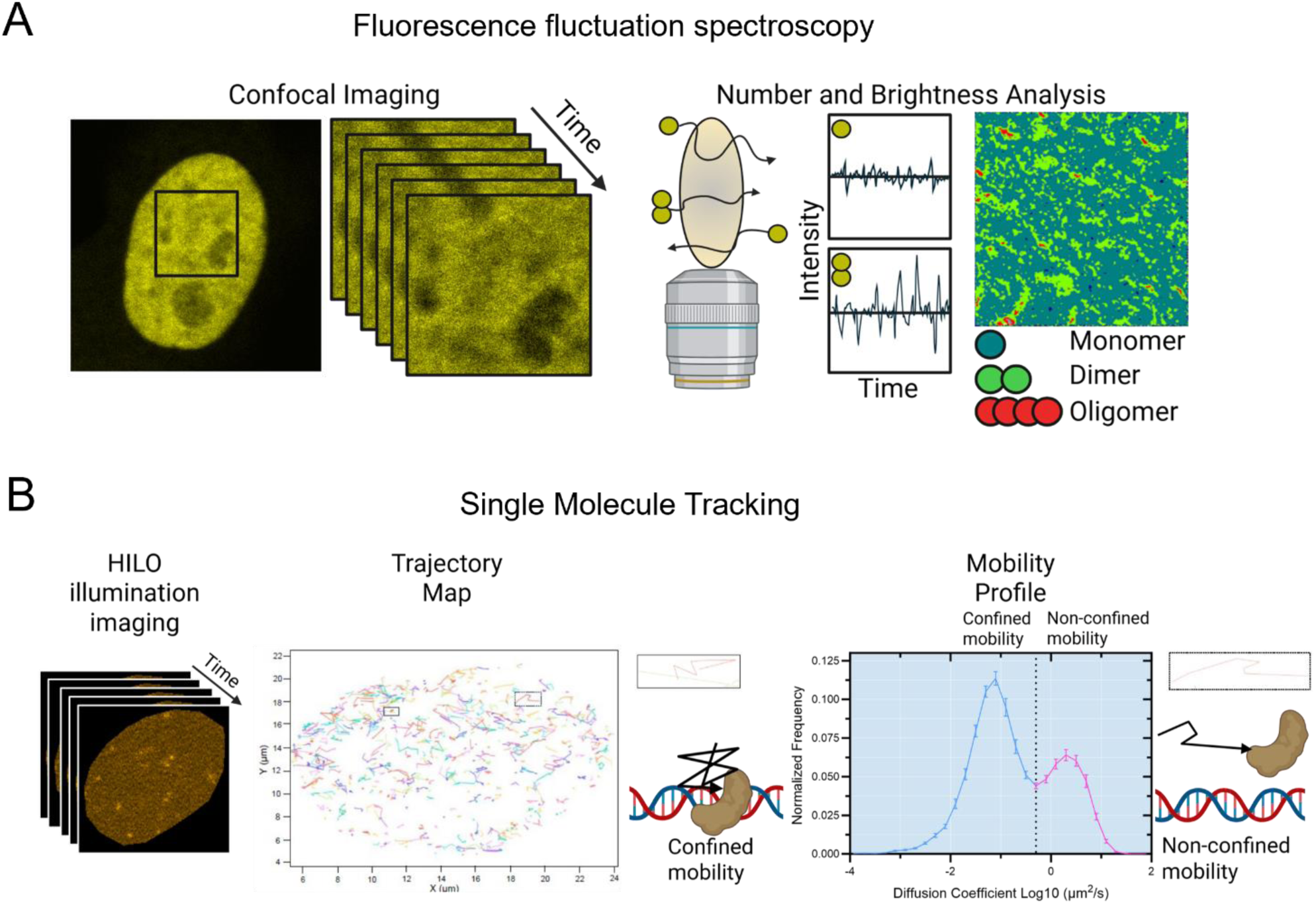
**A.** Number and Brightness (N&B) analysis starts by identifying a cell and raster scanning to generate a confocal time series (100 frames). As the molecules move through the confocal volume the different oligomeric states will cause differences in fluorescence intensity. The intensity fluctuations are assessed over time and converted to fluctuations in molecular brightness for every pixel. The brightness is indicative of the average oligomeric state. In this way a dimer is twice as bright as a monomer and higher order oligomer are brighter than a dimer. **B.** Single molecule tracking (SMT) analysis starts by identifying a cell and imaging by highly inclined and laminated optical sheet (HILO) illumination to generate a time series (6000 frames). From the time series each Halo-tagged SOX18 molecule is identified per frame and stitched together to build a trajectory map. From each trajectory the diffusion coefficient is calculated as a measure of molecular mobility. Trajectories that have a low diffusion coefficient are defined as having a confined mobility (blue), whereas trajectories that have a higher diffusion coefficient are defined as being diffusive (non-confined mobility; pink). The diffusion coefficients are then graphed to assess the proportion of molecular populations that falls into either category.

**Table S1.** Summary of biophysical experiments and biological interpretations.

| SOX18 mobility and oligomeric state is effected by chromatin organization |  |  |  |
| --- | --- | --- | --- |
| N&B (=PPI, oligomeric status) | Biophysical Observation | Biological interpretation | Fig |
| ActD treatment relative to DMSO | No significant difference in the percentage of monomers and dimers, significant increase in oligomers | ActD treatment = loss of chromatin accessibility, leading to the loss of binding locations and subsequent decline in dimer formation. | Fig 3A-D |
| TSA treatment relative to DMSO | Significant increase in percentage of monomers & significant decrease in percentage of dimers, no significant difference in percentage of oligomers | TSA treatment = increase in chromatin accessibility, leading to increased binding locations and an increase in the dimer and higher order oligomer formation | Fig 3A-D |
| **SMT (Diffusion)** | **Biophysical Observation** | **Biological interpretation** | **Fig** |
| TSA treatment relative to DMSO: non-confined mobility (= target search pattern) | Significantly decreases | Increased chromatin accessibility from TSA = more SOX18 molecules binding and less diffusing | Fig 3E-F |
| TSA treatment relative to DMSO: confined mobility (= PPI or protein-chromatin interaction) | Significantly increases |  | Fig 3E-F |
| **SMT (Temporal occupancy)** | **Biophysical Observation** | **Biological interpretation** | **Fig** |
| TSA treatment relative to DMSO: short occupancy (= searching behavior) | No significant differences | Increased chromatin accessibility from TSA = SOX18 forming more dimers and higher order oligomers | Fig 3G-I |
| TSA treatment relative to DMSO: long occupancy (= PPI or protein-chromatin interaction) | Significantly increases |  | Fig 3G-I |
| TSA treatment relative to DMSO: ratio of occupancy | No significant differences |  | Fig 3G-I |

**Supplementary Figure 4. Related to Fig 4.**
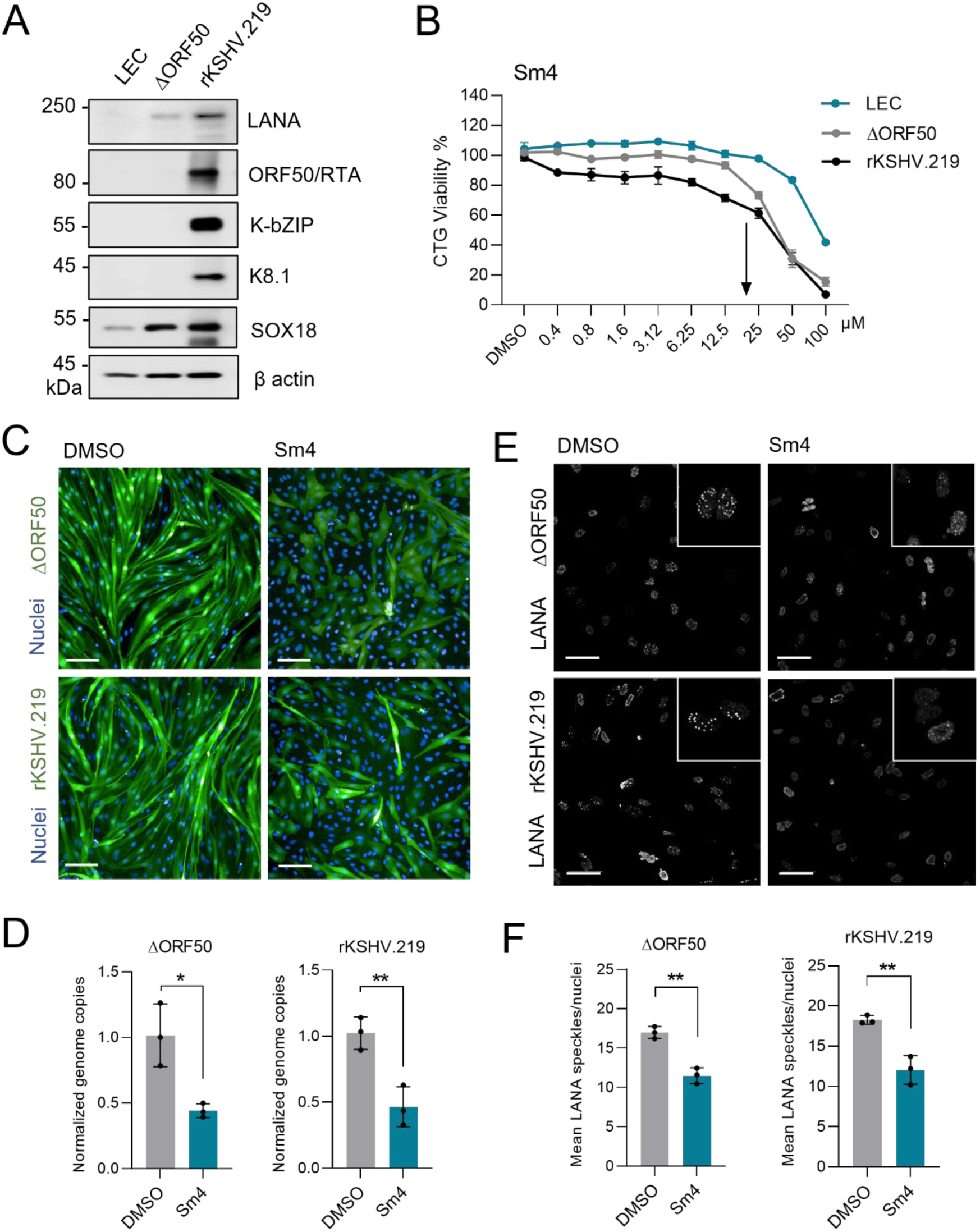
**A.** LECs infected with latent KSHV-BAC16-ΔORF50 (ΔORF50-KLEC) or wildtype rKSHV.219 (rKSHV.219) for 72h and immunoblotted for the indicated viral proteins, and SOX18, using β-actin as a loading control. **B.** CTG viability assay of LECs infected with ΔORF50 or rKSHV.219 and treated with DMSO or with the indicated increasing Sm4 concentrations. **C-F.** Infection phenotypes of LECs infected with GFP-expressing ΔORF50-KLEC or rKSHV.219 and treated at 72h.p.i with Sm4 or DMSO control for 72h. **C.** GFP images of infected cells upon DMSO or Sm4 treatments. Nuclei were counterstained with Hoechst (33342), scale bar is 100µm. **D.** Relative KSHV DNA genome copies. **E.** Images of anti-LANA labeled infected cells and F) quantified as mean from 10 fields for each n=3 biological replicates. Nuclei were counterstained with Hoechst (33342), scale bar is 50µm. Statistical significance was determined by unpaired t-test, *p < 0.05, **p < 0.01.

**Supplementary Figure 5. Related to Fig 4.**
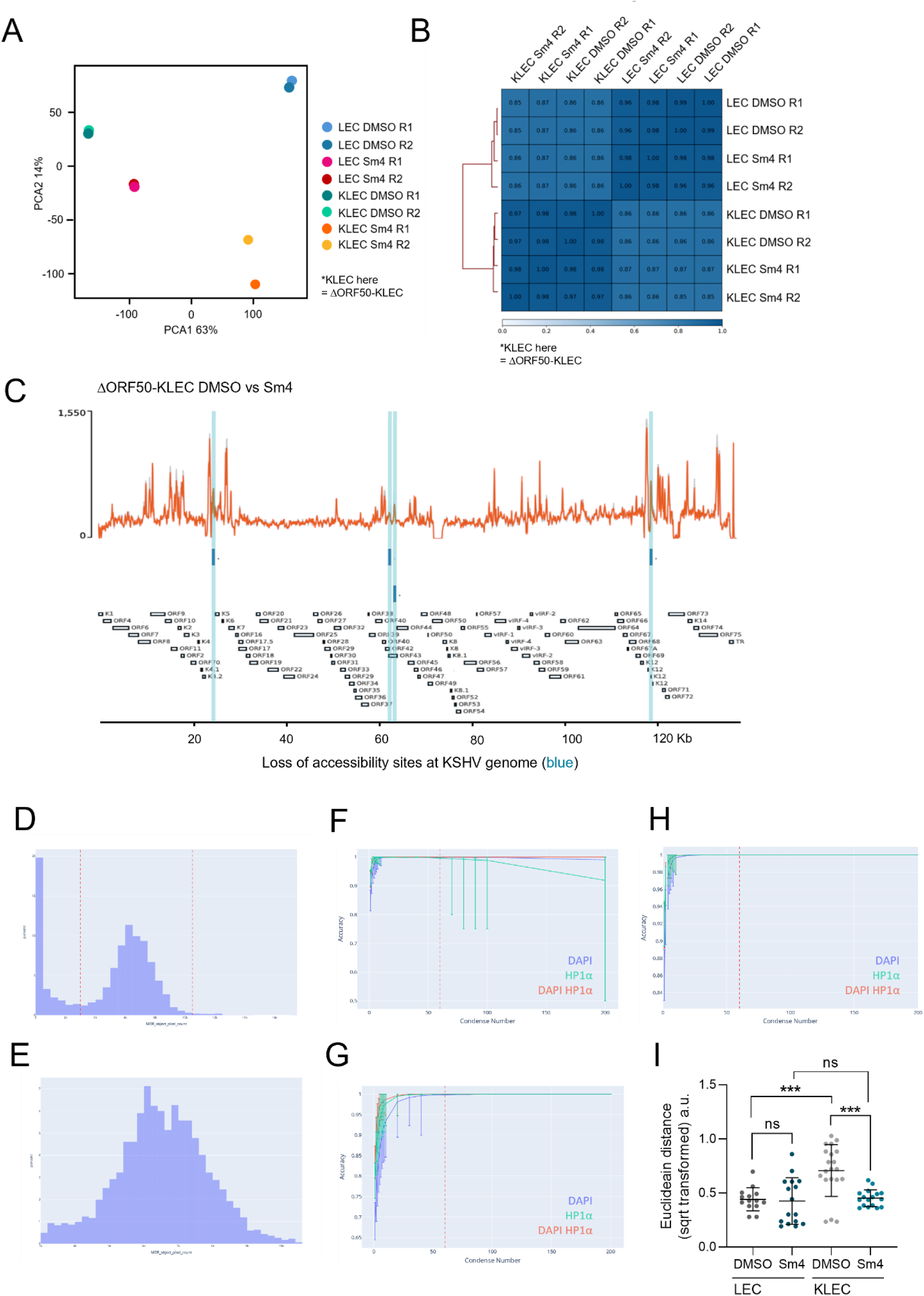
**A-C.** LECs infected with KSHV-BAC16-ΔORF50 (ΔORF50-KLEC) and treated with Sm4 or DMSO for 24h and processed for ATAC-seq. **A.** Clustering and PCA analysis of the ATAC-seq data. **B.** Pearson’s analysis of the replicate samples. **C.** Analysis of the ATAC-seq peaks on the KSHV genome in ΔORF50-KLECs treated with DMSO (grey) or Sm4 (red) indicating loss of accessibility sites (blue). **D.** Histogram displaying distribution of all object sizes (pixels) identified during segmentation, objects inside of red-dashed lines are used in analysis and **E.** zoom in on dashed lines in panel D. **F-H.** LEC and KLEC stained with DAPI and anti-HP1α antibody. Line plots showing accuracy measurements versus cell condense number, 95% confidence intervals are shown. **F.** Cell condensation between LEC and KLEC DMSO treatment. **G.** Cell condensation between LEC DMSO vs Sm4 treatment. **H.** Cell condensation between KLEC DMOS vs SM4 treatment **I.** Euclidean distances square root transformed of points from Fig 4L. n = 14 points or more. Statistical significance was determined by one-way ANOVA with Tukey correction for multiple comparisons, ***p < 0.001, ns = non-significant.

**Supplementary Figure 6. Related to Fig 5.**
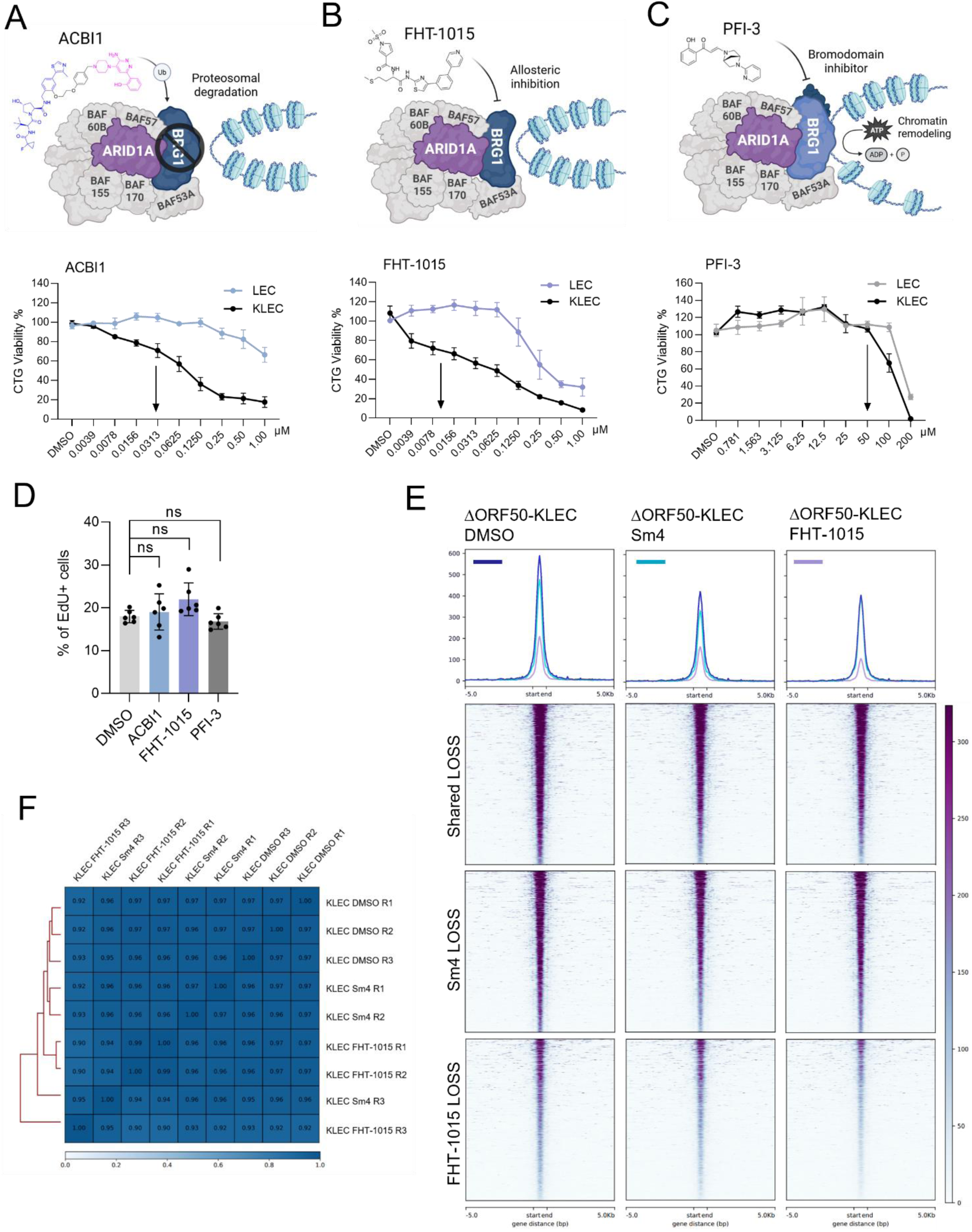
**A-C.** A schematic of the inhibitor mode of action is shown in the top panels. CTG viability assay of uninfected LECs (LEC) or LECs infected with rKSHV.219 (KLEC) for 72h and treated with the indicated, increasing concentrations of BRG1 inhibitors A) ACBI1, B) FHT-1015 and C) PFI-3 (n=3), arrows indicate the selected concentration for following inhibitor assays. **D.** LECs and KLECs were treated with ACBI1, FHT-1015 and PFI-3 and treated with EdU for 4h before subjecting to EdU Click-It, imaged with Opera Phenix 20x and quantified from (n=6 independent replicates, and from each n=100 nuclei). Statistical significance was determined by one-way ANOVA with Dunnett correction for multiple comparisons, ns = non-significant. **E-F.** LECs infected with ΔORF50 and treated with Sm4, FHT-1015, or DMSO for 72h and processed for ATAC-seq. **E.** Peaks and heatmaps of the top 1000 genomic regions with reduced overall accessibility (dark blue maps) showing shared (dark blue line), unique to Sm4 (turquoise) and unique to FHT-1015 (purple) loss sites. **F.** Pearson’s analysis of the replicate (n = 3) samples. ns = non-significant.

**Supplementary Figure 7. Related to Fig 6.**
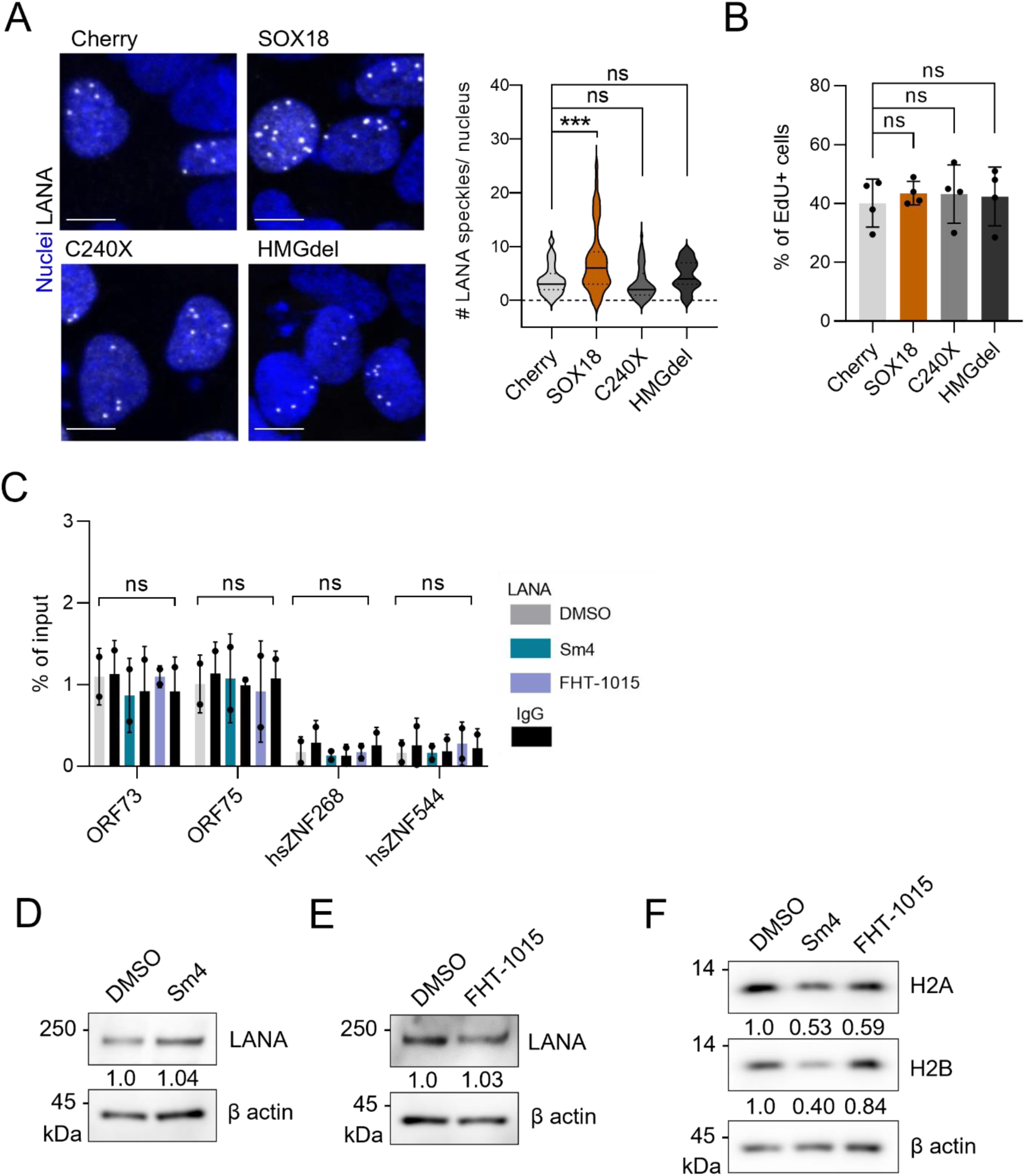
**A.** Confocal images of LANA speckles (above panel) and quantified (bottom panel) as a number of nuclear speckles (n=50 nuclei), nuclei were counterstained with Hoechst (33342), scale bar is 10µm. **B.** HeLa cells expressing SOX18wt or the indicated mutants treated with EdU for 2h before fixing and subjected to EdU Click-It, imaged and quantified (n=4 independent replicates, and from each n=100 nuclei). **C-E.** LECs infected with rKSHV.219 for 72h were treated with DMSO, Sm4 or FHT-1015 for 24h and C) subjected to ChIP-PCR using anti-LANA and IgG antibodies for viral and human non-LANA binding control sites (n=2), and D-E) immunoblotted for LANA and β-actin as a loading control for normalization. **F.** KLECs treated for 72h and immunoblotted for H2A and H2B and quantified as in D-E. Statistical significance was determined by one-way ANOVA with either Dunnett or Tukey correction for multiple comparisons, ***p < 0.001, ns = non-significant.

